# The atypical IκB factor IκBδ enhances CD8 T cell accumulation and effector functions in solid tumors

**DOI:** 10.64898/2026.06.17.732005

**Authors:** Barsha Dash, Xiaocui He, Arlet Lara-Custodio, Leo J Arteaga-Vazquez, Yue Zhao, Hannah Drum, Osamu Ikeda, Eric Johnson, Yuntian Zhu, Chen Zhang, Srikanth Battu, Edahí González-Avalos, Anjana Rao, Patrick G Hogan

**Author notes:** Equal contributions (Barsha Dash, Xiaocui He). Corresponding authors Contact information: Patrick G Hogan, Barsha Dash.

## Abstract

Two prominent mechanisms by which tumors fend off immune control are by constraining the ability of T cells and CAR T cells to survive and expand in the tumor, and by restraining their ability to sustain full cytotoxic capacity. We identified IκBδ, encoded by *Nfkbid*, a poorly characterized IκB family member, as a molecular lever that overcomes both of these constraints on anti-tumor CD8^+^ tumor-infiltrating lymphocytes (TILs). *Nfkbid* is an NFAT target gene that is expressed in CD8^+^ effector T cells and, at modest levels, in CD8^+^ TILs. We found that *Nfkbid* depletion impaired TIL accumulation, exacerbating the growth of solid tumors. On the other hand, ectopic IκBδ overexpression enhanced TIL expansion, reduced the expression of exhaustion-associated transcription factors and inhibitory receptors, and elevated cytotoxic molecule production, leading to enhanced tumor control. IκBδ has a shorter protein isoform that is identical in a core region spanning the ankyrin-repeat domain known to interact with NFκB proteins, but that lacks the ∼150-residue N-terminal region. We showed that the shared core region is sufficient to drive T cell accumulation, whereas the N-terminal peptide region is required for robust effector function and to counter exhaustion, underscoring that tumor-infiltrating CD8^+^ T cell accumulation and effector differentiation are separable programs. Our current study provides evidence that IκBδ, an atypical member of the NFκB family, is a lever to overcome two cardinal deficits that limit CD8^+^ TIL anti-tumor efficacy: impaired accumulation in the tumor and diminished effector function.

## INTRODUCTION

Cancer immunotherapy has underscored the remarkable capacity of T cells to mediate tumor control, yet its clinical success is tempered by progressive ‘exhaustion’ of tumor-infiltrating lymphocytes (TILs) ^1–3^. CD8^+^ T cell exhaustion impairs cytotoxicity directed against tumor cells as well as other effector functions of both endogenous TILs and engineered tumor-specific CAR T cells ^2,3^. Notably, though, CD8⁺ TILs retain open chromatin regions associated with effector programs ^4–6^, raising the question as to whether they can be redirected toward an effector fate by altering the levels of exhaustion-associated and effector-associated transcription factors.

We have focused on crucial transcriptional mechanisms that underlie exhaustion in TILs for many years ^4,7–11^. T cell receptor (TCR) signaling via the transcription factor NFAT initiates CD8^+^ T cell exhaustion ^7^, a counterpoint to the well-characterized role of NFAT in the T cell activation program when it cooperates with AP-1 ^12–15^. Further, TCR signals in tumor antigen-specific TILs elicit the opening of both exhaustion-related and effector-related chromatin regions ^4^. Fully three-quarters of newly opened chromatin sites in the TILs are effector-related, and these effector-related regions are enriched in NFAT-AP1 composite sites and in NFκB and bZIP binding motifs ^4^. Conversely, depletion of exhaustion-associated NR4A-family or TOX-family transcription factors enhances the effector function of CD8^+^ TILs, and correlates with enrichment of open chromatin regions containing NFκB and bZIP binding motifs compared to control TILs ^9,11^. With respect to bZIP signaling, we demonstrated that overexpression of BATF in CD8^+^ TILs attenuates exhaustion and rescues anti-tumor function ^8^, and Lynn e*t al* had similar success with overexpression of c-JUN ^16^. Whether augmenting NFκB signaling can similarly impart anti-tumor properties to TILs has remained an open question.

Here we screened for early targets of NFAT activation in order to gain insight into the ambivalent role of NFAT in both T cell activation and T cell exhaustion. We identified an NFκB modulator, IκBδ (encoded by *Nfkbid*), whose expression is notably dependent on NFAT. IκBδ is an ‘atypical’ IκB factor in the NFκB pathway and is conserved between mice and humans. In contrast to the straightforward inhibitory action of ‘classical’ IκB factors, IκBδ acts as a modulator, and can sway NFκB signaling in both positive and negative directions ^17–20^. The contributions of IκBδ in a tumor setting are unexplored, and this led us to investigate its role in TILs.

We have found, using complementary protein depletion and protein overexpression approaches, that endogenous IκBδ is necessary for the persistence of CD8^+^ TILs in B16-OVA melanoma, an immunologically ‘cold’ tumor. IκBδ depletion sharply decreases tumor-specific OT-I TIL numbers, and nearly completely abolishes the PD1^+^TIM3^+^ TIL subset. Conversely, overexpression of the predominant ∼55 kDa isoform of IκBδ (IκBδ-Long) enables massive OT-I TIL proliferation and accumulation in B16-OVA tumors, as well as anti-CD19 CAR T cell accumulation in MC38-CD19 adenocarcinomas, an immunologically ‘hot’ tumor. Additionally, IκBδ-Long confers enhanced cytotoxicity, counters exhaustion, and confers improved control of tumor burden. Flow cytometry and single-cell RNA-sequencing of pre-transfer CD8^+^ T cells and TILs recovered from tumors at Day 3 and Day 8 after adoptive transfer established that IκBδ-overexpressing TILs differentiate in the tumor to a state akin to that of short-lived effector cells (SLECs) from acute infections. Overexpression of IκBδ-Short— a naturally-occurring isoform of IκBδ that differs from IκBδ-Long solely in the absence of an ∼150-residue N-terminal region— enables TIL proliferation and survival, but not a strong effector response. Thus, TIL survival and TIL effector differentiation in the tumor are under separate T cell-intrinsic controls, both strongly influenced by NFκB signaling.

T cell and CAR T cell adoptive transfer therapies are often characterized as ‘a numbers game’, with the aim being to deliver a sufficient number of T cells with sufficient effector function to gain the upper hand over an established tumor. Our study identifies the atypical nuclear protein IκBδ, its associated signaling pathways, and its downstream targets as new levers that might be exploited to achieve this aim.

## RESULTS

### Calcium signaling and NFAT activation trigger IκBδ expression

To examine immediate transcriptional consequences of NFAT mobilization, we purified total CD8^+^ T cells from spleen and lymph nodes of C57BL/6 mice, activated them for 48 hours with plate-bound anti-CD3 and anti-CD28, and subsequently expanded them for 4 days in the presence of 10 U/mL IL-2 (**Figure 1A**) to promote a memory-like phenotype ^21^. The CD8⁺ T cells were then stimulated *in vitro* with ionomycin for 2, 6, or 18 hours. Ionomycin triggers the release of calcium from ER stores and store-dependent entry of calcium through ORAI plasma-membrane channels ^22,23^, activating the phosphatase calcineurin and causing dephosphorylation and nuclear import of NFAT. Bulk RNA-sequencing (RNA-Seq) (**Figure 1B**) showed that the mRNAs encoding all three NR4A proteins (NR4A1, NR4A2 and NR4A3), NFAT transcriptional targets and key players in CD8⁺ T cell exhaustion, were very strongly induced under these conditions ^7,9^. In addition to the three *Nr4a* genes, the top 50 differentially expressed genes induced by calcium signaling included those encoding 11 other DNA-binding transcription factors, among them *Crem* ^24^ and the *Egr*-family genes *Egr2* and *Egr3* previously associated with exhaustion and T cell tolerance ^25–27^. In keeping with the ambivalent role of calcium-NFAT signaling in T cells^7,28^, genes encoding transcription factors associated with effector differentiation, including *Myc* ^29^*, Irf4* ^30^, and *Irf8* ^31^, were also upregulated (**Fig. 1B**, *left;* **Extended Data 1A**).

**Figure 1:**
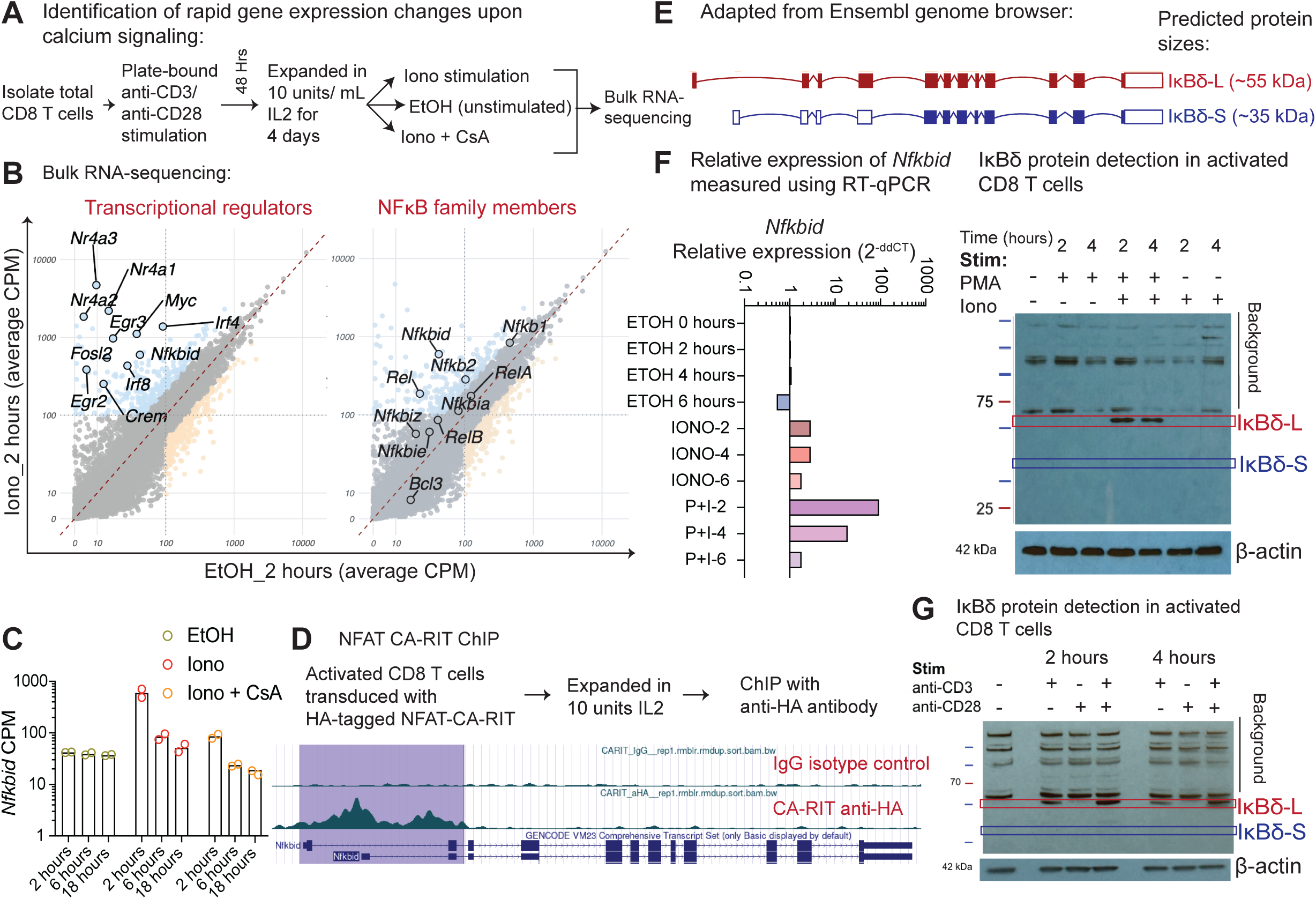
Calcium signaling and NFAT activation trigger IκBδ expression. **(A)** Experimental schematic to identify calcium signaling/NFAT-induced gene expression changes in primary CD8 T cells. CD8 T cells were purified and then activated with plate-bound α-CD3/α-CD28, expanded in 10 units/mL IL-2, and then treated with ionomycin (Iono) or EtOH (vehicle) ± cyclosporin A (CsA), followed by bulk RNA-sequencing. **(B)** Scatterplots showing genes significantly upregulated after 2 hours of ionomycin stimulation compared with EtOH control (purple; cpm >100, adjusted p value < 0.05 and log2 fold-change ≥ 1 computed by DESeq2), including *Nfkbid* (left). Expression of NFκB family member-encoding genes is highlighted (right). Data shown are averaged CPM values from two replicates from one of two independent experiments. **(C)** *Nfkbid* expression (counts per million, CPM) at 2, 6, and 18 h following treatment with EtOH, ionomycin, or ionomycin + CsA. Data shown are two replicates from one of two samples from that underwent library prep on the same day for bulk-RNA sequencing. **(D)** NFAT CA-RIT chromatin immunoprecipitation (ChIP) signal at the *Nfkbid* locus compared with IgG isotype control in activated/expanded CD8 T cells. **(E)** Ensembl genome browser view of two annotated *Nfkbid* splice variants encoding predicted IκBδ isoforms: IκBδ-Long (∼55 kDa) and IκBδ-Short (∼35 kDa). **(F)** Quantitative RT-qPCR was performed to measure relative *Nfkbid* expression in unstimulated, ionomycin treated or PMA and ionomycin treated conditions at 2, 4 and 6 hours (*left*). Expression values were normalized to the housekeeping gene *Rpl13* using the ΔCq method (ΔCq = Cq*_Nfkbid_* – Cq*_Rpl13_*). Relative expression was calculated using the ΔΔCq method with 0-hour ethanol (EtOH) control sample set to 1 (2^−ΔΔCq^). Data shown are the mean of three technical replicates per condition (second independent experiment in supplementary data 1B). IκBδ protein detection in activated CD8 T cells expanded in IL-2. Cell lysates from unstimulated cells and cells treated with PMA, ionomycin, or PMA + ionomycin for the indicated times were analyzed by western blot using a polyclonal α-IκBδ antibody (*right*). Data are representative of three independent experiments. β-actin is the loading control. **(G)** Western blot of CD8 T cells stimulated with α-CD3 and/or α-CD28 for 2- or 4-hours showing induction of IκBδ-Long (L) and IκBδ-Short (S) (right); β-actin is the loading control. Data are representative of two independent experiments.

We focused our attention on *Nfkbid* which was induced by ionomycin stimulation and was among the top 50 upregulated genes (**Figure 1B, Extended Data 1A**). Compared to unstimulated CD8 T cells (EtOH vehicle), cells stimulated with ionomycin for 2 hours increased *Nfkbid* messenger RNA (mRNA) by approximately 14-fold (**Figure 1B, Extended Data 1A, B)**. Treatment with cyclosporin A (CsA), an inhibitor of calcineurin, attenuated *Nfkbid* induction by ∼10-fold relative to ionomycin alone (**Figure 1C, Extended Data 1B**).

*Nfkbid* mRNA levels peaked at 2 hours and declined sharply by 6 hours of ionomycin stimulation, but remained elevated above levels in unstimulated cells and remained sensitive to CsA (**Figure 1C**). Among NF-κB-related genes ^32^, *Rel* (encoding c-Rel) and *Nfkb2* (encoding NFκB p100/p52) transcripts increased by ∼8-fold and ∼3-fold respectively (**Figure 1B**, *right*). *Nfkb1* (encoding *NF-κB* p105/p50)*, RelA* (encoding NF-κB p65)*, RelB, Nfkbia* (encoding IκBα), and the atypical IκB-family genes *Nfkbie* (encoding IκBε), *Nfkbiz* (encoding IκB(), and *Bcl3* exhibited more modest changes or were expressed only at low levels.

The observed blockade of *Nfkbid* mRNA induction by CsA suggested that NFAT targets *Nfkbid*. To confirm NFAT binding to the *Nfkbid* locus, we expressed an HA-tagged constitutively active mutant NFAT in CD8^+^ T cells that had been expanded in 10 U/mL IL-2, and performed chromatin immunoprecipitation and sequencing (ChIP-seq) with an anti-HA antibody. We mapped broad peaks of NFAT binding at the first intron of *Nfkbid* (**Figure 1D**). The findings are consistent with our previously published ChIP-seq data for wild-type NFAT in CD8^+^ T cells that had been expanded in 100 U/ml IL-2 ^7^. In this previously published study, endogenous wild-type NFAT showed no detectable binding in unstimulated cells, as expected because NFAT is cytoplasmic in unstimulated T cells, but exhibited clear enrichment extending across the first intron after 1 hour of PMA and ionomycin stimulation ^7^ (data not shown).

Next, we determined that converging signaling pathways other than calcium signaling matter for *Nfkbid* expression. At the mRNA level, relative to unstimulated CD8⁺ T cells, treatment with PMA and ionomycin together induced higher *Nfkbid* expression compared to ionomycin alone at 2 and 4 hours (**Figure 1F**, *left;* **Extended Data 1C** showing two independent experiments conducted on different days). At the protein level, incubation of CD8⁺ T cells with PMA and ionomycin together induced robust expression of the ∼55-kDa isoform IκBδ-L, whereas stimulation with either PMA alone or ionomycin alone did not yield IκBδ-S (Ensembl annotated isoforms of IκBδ shown in **Figure 1E**) protein detectable by western blot (**Figure 1F**, *right*). Stimulation of memory-like CD8⁺ T cells with anti-CD3 induced detectable IκBδ-L, and anti-CD3 plus anti-CD28 further enhanced IκBδ-L expression (**Figure 1G**). The ∼35-kDa IκBδ-S variant was undetectable under our specific conditions, but both IκBδ-L and IκBδ-S can be expressed simultaneously in Tregs^19^ and conventional CD4^+^ T cells^33^. We found conservation between mouse and human IκBδ proteins (**Extended Data 1D**), with ∼80% identity across the full-length protein. Analysis of publicly available human datasets (**DICE database**) demonstrated that *Nfkbid* is also induced upon activation of human CD8⁺ T cells indicating its function is likely similar in mouse and human T cells (**Extended Data 1E**). Further experimental dissection of the pathways that control IκBδ expression will be important for a complete understanding of the roles of IκBδ in cell biology.

### IκBδ is required for accumulation of tumor infiltrating lymphocytes

We noted modest *Nfkbid* expression in exhausted T cells during a chronic infection in a previously published investigation from our group, suggesting a potential role in T cell exhaustion ^34^ (**Extended Data 2A**). To examine functional roles of *Nfkbid in vivo*, we designed a CRISPR (cr) RNA targeting the 6^th^ exon of *Nfkbid*, a region common to both the long and short isoforms (**Figure 2A**, *top*). The crRNA was complexed with Cas9 protein and tracrRNA, and electroporated into activated OT-I CD8⁺ T cells, followed by expansion of the cells in standard T cell medium containing 10 U/ml IL2. Stimulation of *sgNfkbid* treated T cells with PMA and ionomycin yielded only faint IκBδ-L protein bands at 2 hours and 4 hours post-stimulation compared to control sgRNA-treated cells, confirming depletion (knockout, KO) in a majority of cells (**Figure 2A**, *bottom*). After restimulation with PMA and ionomycin, IκBδ KO and control CD8^+^ T cells showed comparable upregulation of CD69 and NFAT2, two key markers of early T cell activation, indicating that the failure of *sgNfkbid*-treated cells to upregulate IκBδ-L upon restimulation was not due to defects in reactivation but rather due to successful *Nfkbid* disruption in most cells (**Extended Data 2B**).

**Figure 2:**
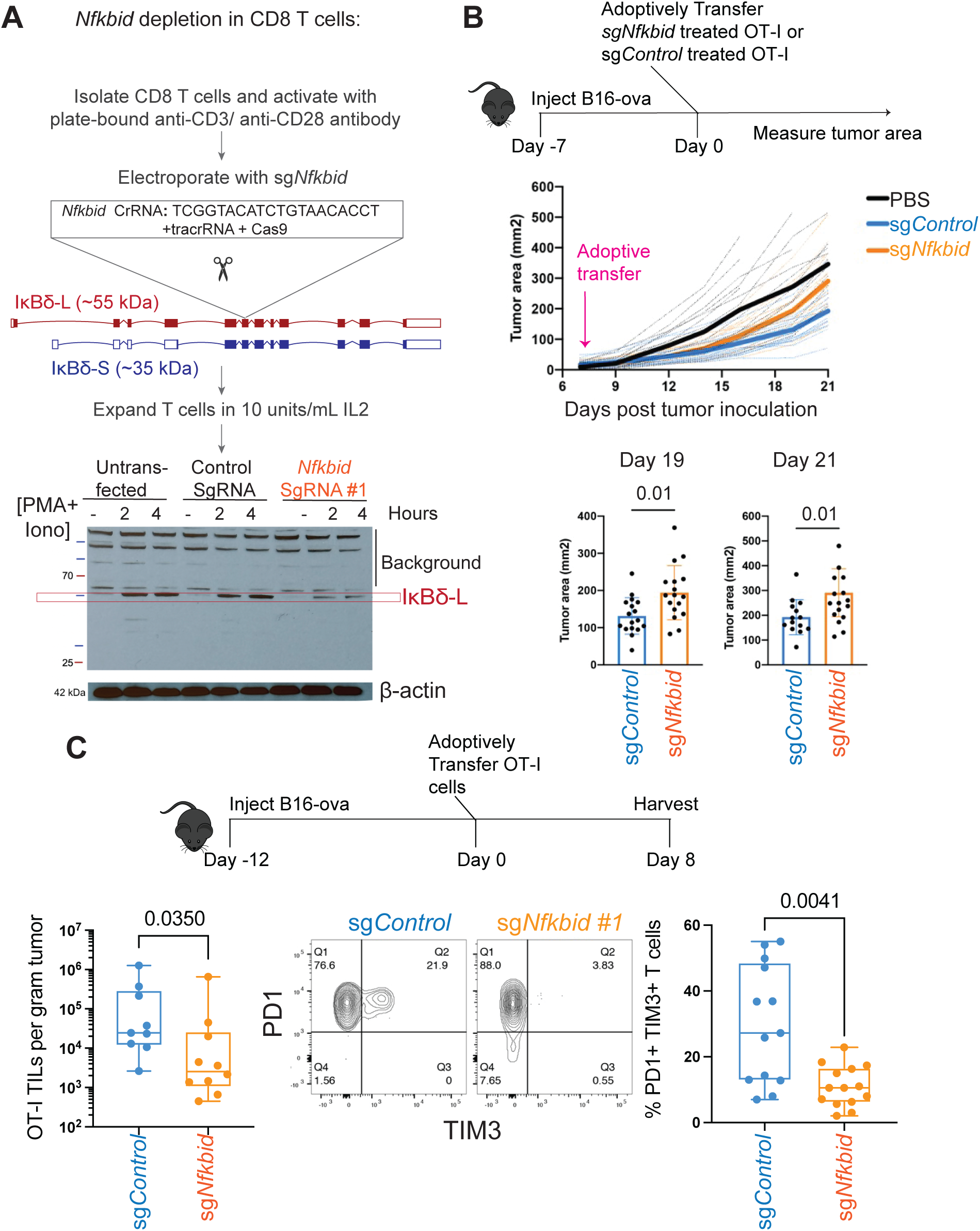
IκBδ is required for accumulation of tumor infiltrating lymphocytes. **(A)** Schematic of CRISPR–Cas9–mediated *Nfkbid* deletion in primary CD8 T cells. CD8 T cells were isolated and activated with plate-bound α-CD3/α-CD28 antibodies, electroporated with Cas9 ribonucleoprotein complexes containing either *Nfkbid*-targeting sgRNA or control sgRNA, and expanded for 4 days in 10 U/mL IL-2. Targeting of the *Nfkbid* locus disrupts both IκBδ-Long (L) and IκBδ-Short (S) isoforms. Efficient depletion of IκBδ-L protein was confirmed by western blot following PMA/ionomycin stimulation; β-actin is the loading control. **(B)** Experimental timeline for adoptive T cell transfer into tumor-bearing mice. B16-OVA melanoma cells were injected subcutaneously, followed by adoptive transfer of OT-I CD8 T cells treated with control sgRNA (sgControl) or *Nfkbid* sgRNA (Sg*Nfkbid*). Tumor growth was monitored longitudinally. Tumor area measurements over time are shown (*top*), with individual tumor areas quantified at days 19 and 21 post-tumor inoculation (*bottom*). Statistical significance was determined using two sample t-test. **(C)** Experimental schematic and analysis of tumor-infiltrating OT-I CD8 T cells. Mice bearing established B16-OVA tumors received adoptive transfer of control or *Nfkbid*-deficient OT-I T cells, and tumors were harvested on day 8 after adoptive transfer. *Left*, OT-I tumor-infiltrating lymphocyte (TIL) counts normalized to mass of tumor (sgControl n=9, and sg*Nfkbid* n=10; from three independent replicate experiments and 19 mice). One of the replicates tested two distinct sg*Nfkbid* reagents (sg*Nfkbid*#1 n=2, and sg*Nfkbid*#2 n=2). *Center*, Representative flow cytometry plots from the same experimental series, showing PD-1 and Tim-3 expression within OT-I CD8 T cells. Data shown are for sgControl and sg*Nfkbid*#1. sg*Nfkbid*#2 had a comparable effect. *Right*, Quantification of PD-1^+^Tim-3^+^ exhausted CD8 T cells (sgControl n=14, and sg*Nfkbid* n=16) with data added from a fourth independent replicate experiment (sgControl n=4 and sg*Nfkbid*#2 n=4; from 8 mice). Statistical significance was determined using two-tailed Mann-Whitney test.

To determine the effects of *Nfkbid* depletion *in vivo*, we examined tumor growth in mice bearing B16-OVA melanoma tumors. 1.5 million control or *sgNfkbid* OT-I CD8 T cells, which recognize an ovalbumin peptide presented by MHC Class I, were adoptively transferred into C57BL/6 mice bearing incipient 7-day-old B16-OVA tumors as previously described ^8,9,11^, and tumor areas were measured every other day (**Figure 2B**, *top*). Compared to mice that received no T cells, both sg*Nfkbid* and sg*Control* OT-I recipient mice exhibited a reduced rate of tumor growth from day 10 to day 15 (**Figure 2B**, *middle*). However, the rate of tumor growth accelerated after day 15 in mice that received *sgNfkbid* OT-I cells, and these mice had significantly larger tumors at days 19 and day 21 than mice that received sg*Control* OT-I cells (**Figure 2B**, *bottom*). When sg*Nfkbid* or sg*Control* CD45.1 OT-I cells were transferred into CD45.2 congenic C57BL/6 mice bearing established 12-day-old B16-OVA as described previously^8,9,11^ (**Figure 2C**, *top*), there were markedly fewer *sgNfkbid* OT-I TILs compared to control TILs at day 8 post adoptive transfer (**Figure 2C**, *bottom left*). The frequency of PD1⁺Tim3⁺ “effector-exhausted” TILs was conspicuously reduced for sg*Nfkbid* TILs compared to sg*Control* TILs, even after taking into account the lower number of sg*Nfkbid* TILs. The few detected PD1⁺Tim3⁺ cells could be escapees which failed to delete IκBδ. The data indicate that IκBδ expression is central to the persistence of OT-I TILs in B16-OVA tumors, to the differentiation or survival of PD1⁺Tim3⁺ TIL subset, and to early tumor control.

### IκBδ-L overexpression enhances TIL proliferation and numbers

Next, we asked whether ectopic expression of IκBδ would elicit effects opposite to IκBδ depletion. We addressed this by overexpressing the predominant protein isoform IκBδ-L in CD8⁺ T cells using an MSCV retroviral vector (pMIG) that also drives the expression of the green fluorescent protein (GFP) reporter (**Figure 3A, Extended Data 3A**). We achieved ∼90% transduction efficiency (**Extended Data 3A**) and confirmed protein expression by western blotting (**Figure 3A**). For TIL experiments, CD45.1⁺ OT-I CD8 T cells were transduced with either an empty vector (EV) or the IκBδ-L expression plasmid (IκBδ-L-OE), and subsequently transferred into congenic CD45.2⁺ C57BL/6 mice bearing 12-day-old established B16-OVA tumors (**Figure 3B**). Tumors were harvested at days 3, 5, and 8, dissociated, and the tumor-infiltrating lymphocytes of interest were identified by CD45.1 and CD8 expression as well as GFP expression by flow cytometry. OT-I TILs expressing IκBδ-L displayed a pronounced increase in absolute numbers relative to EV control TILs on day 5 and day 8 (**Figure 3C**) and an increased frequency compared to control TILs on day 8 (**Figure 3D**). At day 8, mice which received OT-I TILs expressing IκBδ-L exhibited slightly smaller tumors on average and this difference was statistically significant (**Figure 3E**).

**Figure 3:**
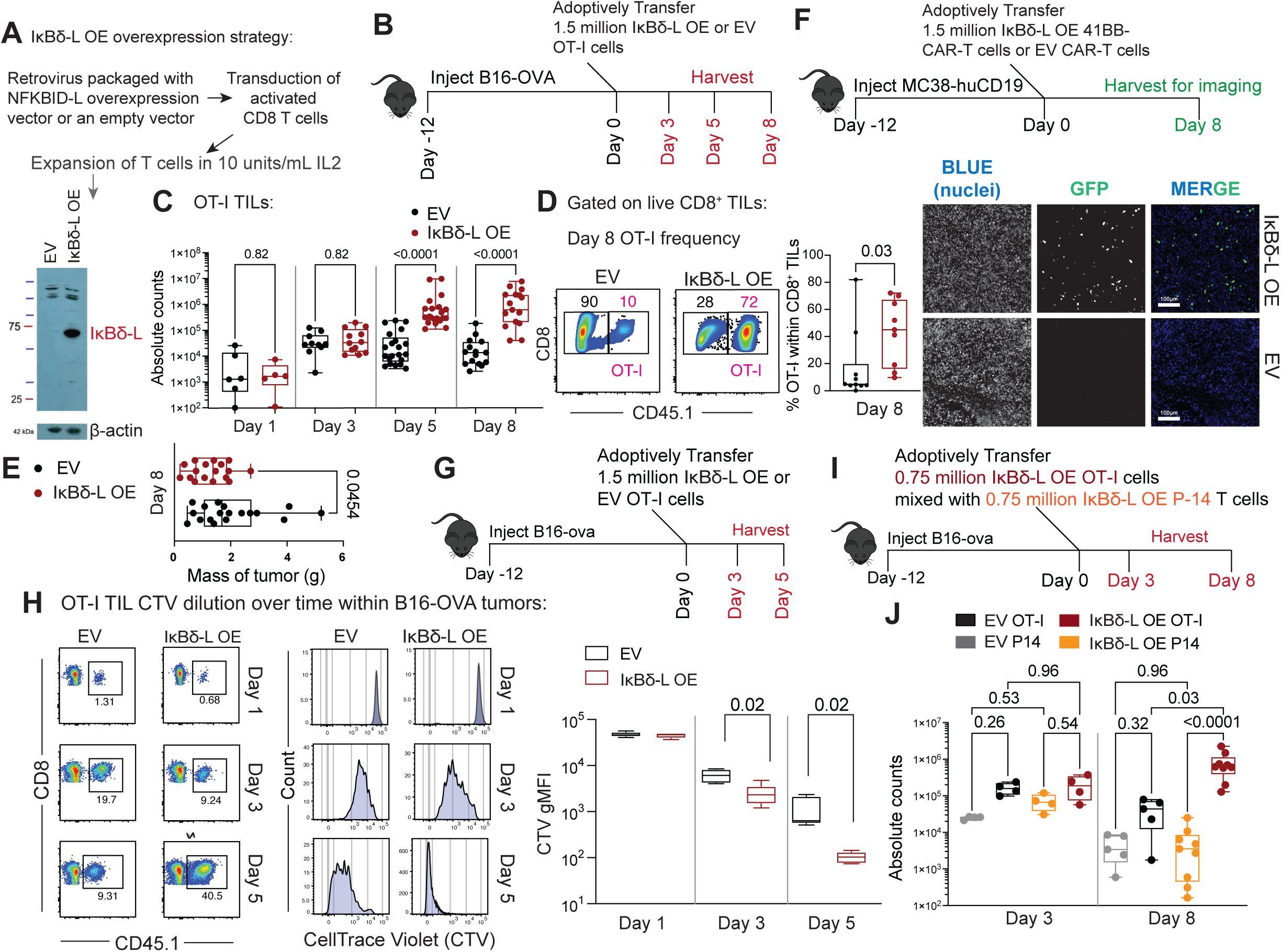
IκBδ-L overexpression enhances TIL proliferation and numbers. **(A)** Strategy for retroviral overexpression of IκBδ-Long (or IκBδ-L) in activated CD8 T cells. Activated CD8 T cells were transduced with retrovirus encoding IκBδ-L or empty vector (EV) control (MSCV backbone) and expanded in 10 units/mL IL-2. Overexpression of IκBδ-L protein was confirmed by western blot; β-actin is the loading control. Data representative of 7 independent experiments. **(B)** Experimental timeline for adoptive transfer of IκBδ-L-overexpressing or EV OT-I CD8 T cells into B16-OVA tumor–bearing mice. Mice received 1.5 × 10⁶ OT-I cells and tumors were harvested at days 3, 5 and 8. **(C)** Absolute numbers of OT-I tumor-infiltrating lymphocytes (TILs) recovered from B16-OVA tumors at days 3, 5, and 8 following adoptive transfer of EV or IκBδ-L-overexpressing OT-I cells. Each point represents an individual mouse with n=5-17 mice per condition pooled from at least 3 independent experiments per time point (Day 1: EV n=6, IκBδ-L n=5; Day 3: EV n=10, IκBδ-L n=12; Day 5: EV n=21, IκBδ-L n=21; Day 8: EV n=15, IκBδ-L n=17). Statistical significance was determined using one-way ANOVA (non-parametric Kruskal-Wallis test with Benjamini and Hochberg correction for multiple comparisons for controlling the False Discovery Rate). **(D)** Representative flow cytometry plots showing OT-I frequency among live CD8⁺ TILs at day 8 following adoptive transfer of EV or IκBδ-L-overexpressing OT-I cells, and quantification of frequency of OT-I cells within total CD8 T cells. Each point represents an individual mouse with n=9-10. Statistical significance was determined using unpaired t-test (Day 8: EV n=10, IκBδ-L n=12). **(F)** Mass of tumors resected from mice that received either EV OT-I or IκBδ-L-overexpressing OT-I T cells at day 8 (EV n=18, IκBδ-L n=17). Data from 4 independent experiments. Statistical significance was determined using unpaired t-test. **(F)** Experimental design for evaluating the effect of IκBδ-L overexpression on 41BB-CAR-T cell accumulation within MC38-huCD19 tumors. MC38-huCD19 tumor-bearing mice received 1.5 × 10^6^ empty vector (EV) or IκBδ-L-overexpressing (IκBδ-L OE) 41BB-CAR-T cells, and tumors were harvested for imaging on day 8. Representative fluorescence microscopy images of tumor sections showing nuclei (blue), GFP-expressing transferred CAR-T cells (green), and merged channels. **(G)** Experimental timeline for adoptive transfer of CellTrace Violet (CTV) labeled EV or IκBδ-L-overexpressing OT-I CD8 T cells into B16-OVA tumor–bearing mice, followed by tumor harvest at indicated time points. **(H)** Representative flow cytometry plots and CTV dilution profiles of OT-I CD8⁺ TILs isolated at days 1, 3, and 5 post-transfer (*left*), demonstrating enhanced proliferation of IκBδ-L-overexpressing OT-I cells compared with EV controls. Representative data from two independent experiments. Quantification of CTV geometric mean fluorescence intensity (gMFI) in OT-I CD8⁺ TILs at days 1, 3, and 5 post-transfer (*right*) with n=4-5 per condition per time point in two independent experiments (Day 1: EV n=4, IκBδ-L n=4; Day 3: EV n=4, IκBδ-L n=5, Day 5: EV n=5, IκBδ-L n=5). Statistical significance was determined using unpaired t-test within each time point. Data from two independent experiments. **(I)** Experimental design for competitive adoptive transfer of equal numbers (0.75 × 10⁶ each) of IκBδ-L-overexpressing OT-I cells mixed with IκBδ-L-overexpressing P14 cells into B16-OVA tumor–bearing mice. **(J)** Absolute counts of OT-I and P14 TILs recovered from tumors at days 3 and 8 following adoptive co-transfer. Each symbol represents an individual mouse. Each point represents an individual mouse with n=4-9 mice per condition. Data are from two independent experiments at D8 and one experiment at D3 (Day 3: EV n=4, IκBδ-L n=4, Day 8: EV n=5, IκBδ-L n=9). Statistical significance was determined using one-way ANOVA (non-parametric Kruskal-Wallis test with Benjamini and Hochberg correction for multiple comparisons for controlling the False Discovery Rate).

To validate TIL expansion in a second tumor model, we evaluated TIL accumulation in MC38 colon tumors engineered to express human CD19 (**Figure 3F**). Activated CD8 T cells were co-transduced with an anti-CD19 CAR-41BB and either the EV or IκBδ-L MSCV expression plasmid, and transferred into tumor-bearing mice. Tumors were collected at days 2, 5, and 8 and fixed in PFA, and frozen sections were then analyzed by microscopy. GFP^+^ CAR-TILs were not observed on day 2, and EV CAR-TILs remained relatively sparse at the later times, whereas the density of IκBδ-L OE CAR-TILs per unit area increased greatly on days 5 and 8 (**Figure 3F and Extended Data 3B**). We confirmed enhanced IκBδ-L OE CAR-T cell accumulation compared to EV TILs at day 8 using flow cytometry (**Extended Data 3D,** *top and bottom panels*).

To assess whether the increased numbers of IκBδ-L TILs reflected enhanced proliferation, CellTrace Violet (CTV) labeled IκBδ-L overexpressing OT-I cells or EV OT-I cells were transferred into separate B16-OVA–bearing hosts and analyzed at days 1, 3, and 5 for CTV dilution (**Figure 3G-H**). At day 1, the frequencies of CTV⁺ cells were similar, indicating comparable homing to the tumor, and both EV and IκBδ-L OE OT-I populations showed single CTV peaks of high mean fluorescence intensity (MFI), indicating undivided cells (**Figure 3H**, *top panels*). By day 3, both EV and IκBδ-L OE TILs had divided and diluted the CTV signal, with the IκBδ-L TILs undergoing more divisions (**Figure 3H**, *middle panels*). Both cell populations continued dividing through day 5, with the IκBδ-L TILs diluting the CTV signal to near background by day 8 (**Figure 3G**, *bottom panels;* **Figure 3H**). Notably, in the case of EV TILs, the substantial dilution of CTV was accompanied by a net decrease in cell numbers, indicating impaired survival or retention in the tumor microenvironment.

To determine whether accumulation of IκBδ-L expressing TILs was antigen-dependent, we transferred a 1:1 mixture of IκBδ-L OE CD45.1^+^ OT-I cells and of IκBδ-L OE THY1.1^+^ P14 CD8 T cells that do not recognize the OVA peptide into THY1.2^+^ CD45.2^+^ hosts bearing the same B16-OVA tumors (**Figure 3I**). As control, empty-vector (EV)-transduced OT-I and P14 cells were also mixed in a 1:1 ratio and adoptively transferred into tumor-bearing mice. At day 3, EV OT-I, EV P14 cells and IκBδ-L OE OT-I and P14 cells were all detectable in the tumors (**Figure 3J**, *left*). At day 8, IκBδ-L OE P14 T cells had failed to expand, while IκBδ-L OE OT-I TILs exhibited expansion and persistence within the tumors (**Figure 3J**, *right*). Together these data demonstrate that IκBδ-L expression enhances tumor antigen-specific proliferation and survival in solid tumors.

### IκBδ-L redirects TIL differentiation and tempers expression of exhaustion markers

To parse how the gene expression landscape of IκBδ-L-OE TILs changed at the level of single cells as a function of time in comparison to EV TILs, we performed single-cell RNA sequencing (scRNA-seq) on OT-I cells prior to transfer into tumor-bearing mice, and on tumor-infiltrating lymphocytes (TILs) harvested on days 3 and 8 (**Figure 4A, Extended Data 4A**). Pre-transfer EV and IκBδ-L OE OT-I T cells exhibited largely overlapping distributions on two-dimensional UMAP embeddings. However, by day 3 post-transfer into B16-OVA bearing mice, the two populations began to diverge, and by day 8, EV and IκBδ-L OE TILs largely occupied distinct regions of the UMAP space (**Figure 4B**). Two-dimensional UMAP embeddings of *Tcf7* and *Gzmb* expression revealed that IκBδ-L OE TILs were enriched in clusters characterized by high *Gzmb* expression, whereas EV TILs were preferentially represented in clusters expressing *Tcf7* (**Figure 4C**). Pseudobulk expression of mRNAs encoding cytotoxic molecules, *Gzma*, *Gzmb*, *Gzmk* and *Prf1* was elevated by D8 in IκBδ-L OE TILs compared to EV (**Figure 4D**). Notably, genes encoding killer cell lectin-like receptors ^35^ (*Klrc1*, *Klrc2*, *Klrd1*, and *Klrk1*) showed moderately higher or more sustained expression at D8 in IκBδ-L OE TILs. In contrast, expression of the exhaustion-associated transcription factors *Tox* and *Nr4a1/2/3* was reduced relative to EV TILs. Further pseudobulk analysis at day 3 indicated that EV TILs preferentially expressed other exhaustion-associated genes such as *Nrgn*, whereas IκBδ-L-overexpressing TILs were enriched for cytotoxic effector genes and also *Klf2* consistent with early skewing toward an effector program (**Extended Data 4B**). Compared to D3 and D8 EV TILs, IκBδ-L OE D3 and D8 TILs exhibited enhanced expression of *S100a4* and *S100a6*, modulators of calcium signaling^36^ previously reported to be upregulated in early effector OTI T cells targeting acute infections with VSV-OVA or *Listeria*-OVA (**Immgen database,** microarray gene skyline). Compared to D3 and D8 EV TILs, IκBδ-L OE D3 and D8 TILs also exhibited enhanced expression of *Ccl5*, which encodes a chemoattractant for both innate and adaptive immune cells indicating enhanced inflammation (**Extended Data 4C)**.

**Figure 4:**
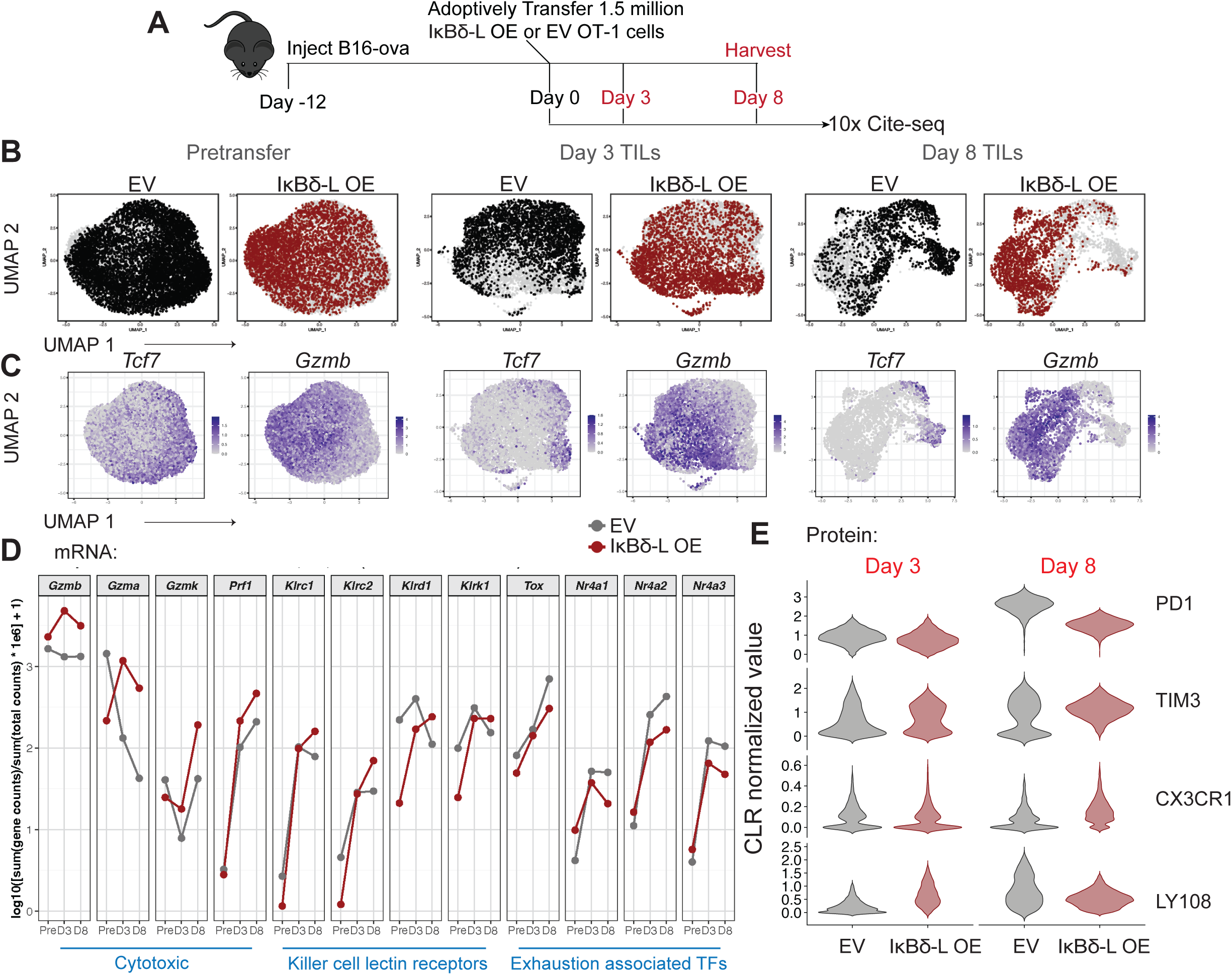
IκBδ-L redirects TIL differentiation and tempers expression of exhaustion markers. **(A)** Experimental schematic for 10x Genomics single-cell RNA-seq with CITE-seq for EV and IκBδ-L overexpressing pre-transfer OT-I cells and OT-I TILs. Mice bearing established B16-OVA tumors received adoptive transfer of 1.5 × 10⁶ OT-I CD8⁺ T cells transduced with empty vector (EV) or IκBδ-L encoding vector. Tumor-infiltrating OT-I lymphocytes (TILs) were harvested at day 3 or day 8 post-transfer and analyzed by 10x Genomics single-cell RNA-seq with CITE-seq. Cells were pooled from n= 3-4 mice per condition per time point *in vivo*. **(B)** UMAP projections of single-cell transcriptomes from pre-transfer OT-I cells, day 3 TILs, and day 8 TILs, colored by EV or IκBδ-L expression. **(C)** Feature plots showing expression of progenitor-associated (*Tcf7*) and effector-associated (*Gzmb*) transcripts across pre-transfer cells and TILs at days 3 and 8. **(D)** Line graphs showing average normalized CPM for effector (*Gzmb*, *Gzma*), Killer cell lectin receptors (*Klrc1, Klrc2, Klrd1, Klrk1*), and exhaustion-associated (*Tox, Nr4a1, Nr4a2, Nr4a3*) in EV and IκBδ-L overexpressing OT-I cells pre-transfer and at days 3 and 8 post-transfer. **(E)** CITE-seq-derived protein expression profiles of tumor-infiltrating OT-I cells. Violin plots show centered log-ratio (CLR) normalized antibody-derived tag (ADT) expression of surface markers (CD8α, TIM-3 [CD366], Ly108, CD62L, PD-1 [CD279], CX3CR1, CD127, and KLRG1) in EV- and IκBδ-L–transduced OT-I T cells recovered from tumors at day 3 and day 8 post-transfer (extreme outliers were defined as values above 99^th^ percentile within each marker and were trimmed for violin plots but are reported in Extended Data 4D as box and whisker plots).

CITE-seq epitope analysis revealed reduced PD1 and TIM3 protein expression in IκBδ-L-OE TILs, particularly by day 8 (**Figure 4E and Extended Data 4D**). Notably, the progenitor marker LY108 was initially higher in IκBδ-L-OE cells, pre-transfer and at day 3, but did not mirror the progressive increase seen in control cells, whereas CX3CR1— ordinarily a marker of CD8^+^ TILs with the highest effector function— increased over time. IL7Rα expression decreased and KLRG1 expression increased in IκBδ-L-OE TILs by day 8, reminiscent of cells with a short-lived effector phenotype found in acute infections (**Extended Data 4D**) Together, these data suggest that IκBδ-L drives a transcriptional and phenotypic shift from a progenitor like state toward a more differentiated cytotoxic effector program in tumor infiltrating CD8 T cells.

### IκBδ-L regulates the speed of progenitor differentiation into effector TILs

Unsupervised clustering of day 8 tumor-infiltrating lymphocytes (TILs) identified seven transcriptionally distinct populations (clusters 0-6) (**Figure 5A**). We have placed them in the order 3, 4, 5, 2, 6, 0, 1 based on decreasing *Tcf7* expression, although the ordering of the last three on this basis is statistically uncertain, given their low expression of *Tcf7* (the ordering is for practical presentation purposes, and is not intended to imply a developmental progression). Clusters 3 and 4 were characterized by high *Tcf7* expression, cluster 5 stood out for elevated expression of the proliferation marker *Mki67*, and clusters 2, 6, 0, and 1 were enriched for *Gzmb* (**Figure 5B**). IκBδ-L overexpression altered the distribution of TILs across these states, with IκBδ-L OE TILs preferentially found in the *Gzmb^high^* effector-like clusters, whereas EV TILs were more uniformly distributed in the *Tcf7^high^* progenitor-like and *Gzmb^high^* effector-like clusters (**Figure 5C**) Despite this shift in proportions, predicted absolute cell numbers, estimated by combining cluster frequencies with total TIL counts from Figure 3 and presented here on a logarithmic scale, were increased across all clusters in IκBδ-L OE TILs (**Figure 5C**). IκBδ-L expands the overall T cell compartment while skewing differentiation toward effector-like states. Flow cytometric analysis confirmed that although the frequency of TCF1⁺ OT-I TILs was reduced for IκBδ-L OE TILs, the absolute number of TCF1⁺ TILs was significantly increased **(Figure 5D**), that is, progenitor-like cells were numerically expanded despite their reduced proportional representation. We examined the expression of selected genes in clusters that correspond to the progenitor-exhausted (clusters 3-4), rapidly proliferating (cluster 5), and effector-exhausted (clusters 2, 6, 0, 1) states of wild-type CD8^+^ TILs (**Figure 5E, Extended Data 5A**). The analysis revealed a continuum in which progenitor-associated genes *(Bcl6* ^37^*, Ccr7*) were downregulated in concert with *Tcf7*, while cytotoxic and effector-associated genes (*Prdm1*, *Gzmb, Prf1, Cxcr6*) were upregulated (**Figure 5E and Extended Data 5A**). Notably, IκBδ-L overexpression led to premature loss of *Bcl6*, *Nr4a1*, and *Nrgn* in the *Tcf7*^+^ clusters, and concomitant early upregulation of *Gzma*, *Gzmb*, *Gzmk*, *Prf1*, and *Klf2* ^38^ in these clusters. In the proliferative cluster 5, IκBδ-L expressing TILs exhibited greater expression of *Gzma/b/k*, *Mki67* and reduced expression of *Nrgn* and *Ccr7* (**Extended Data 5A**).

**Figure 5:**
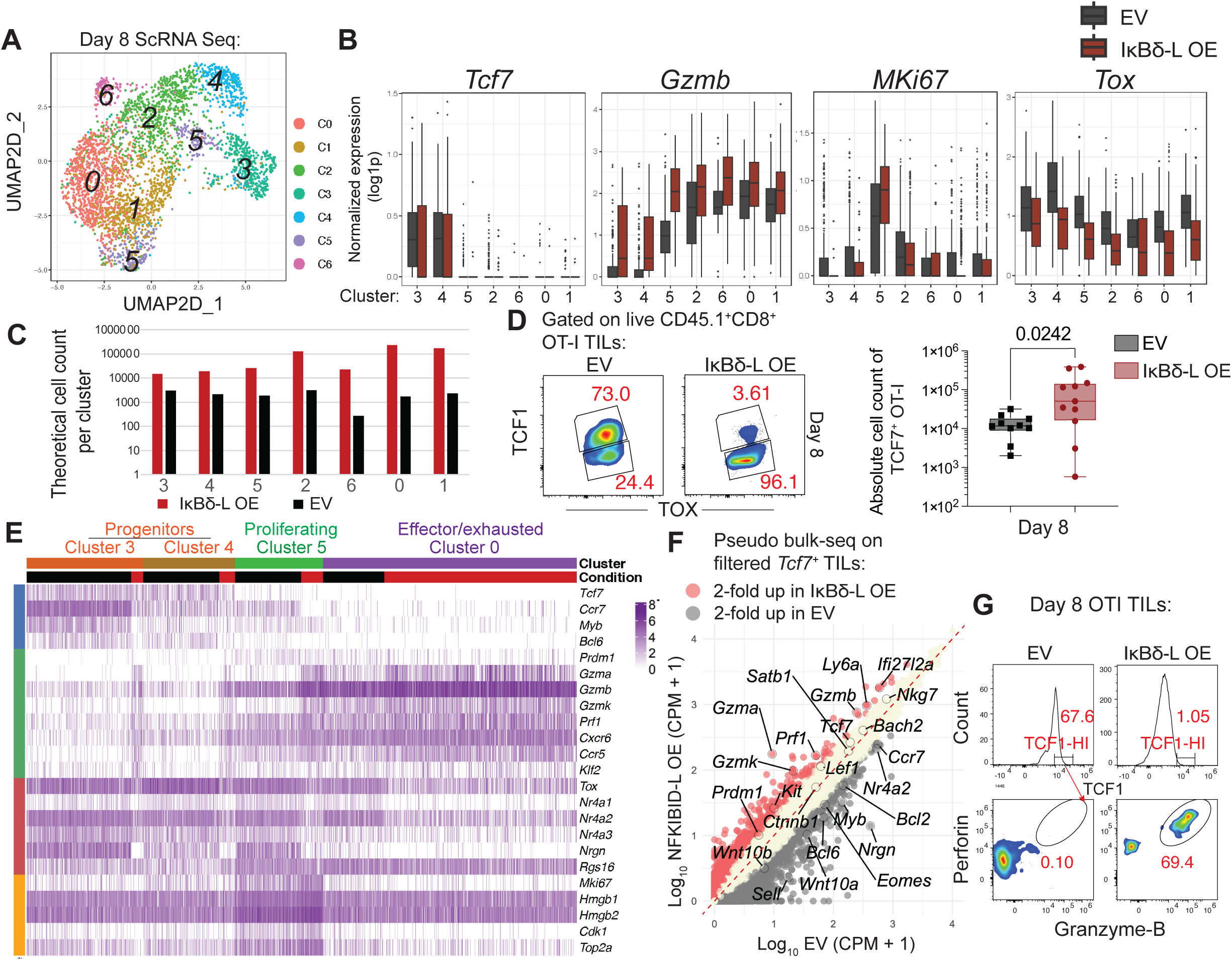
IκBδ-L regulates the speed of progenitor differentiation into effector TILs. **(A)** UMAP projection of day 8 tumor-infiltrating OT-I CD8⁺ T cells analyzed by single-cell RNA-sequencing (scRNA-seq). Cells are grouped into transcriptionally distinct clusters (resolution = 0.5) identified from EV and IκBδ-L–overexpressing (OE) populations. **(B)** Boxplots showing normalized gene expression across clusters for progenitor-associated (*Tcf7*), effector (*Gzmb*), proliferation (*Mki67*), and exhaustion (*Tox*) transcripts comparing EV and IκBδ-L OE OT-I TILs at day 8. **(C)** Theoretical counts of EV and IκBδ-L OE cells across transcriptional clusters, illustrating enrichment of IκBδ-L OE cells within all clusters. **(D)** Representative flow cytometry plots of TCF1 and TOX expression in live CD45.1⁺CD8⁺ OT-I TILs at day 8 following adoptive transfer of EV or IκBδ-L OE cells (*top*). Quantification of absolute numbers of TCF7⁺ OT-I TILs recovered from tumors (*bottom*). Each point represents an individual mouse; statistical significance was determined using an unpaired t-test and data are representative of three experiments. **(E)** Heatmap showing normalized expression of selected genes across four out of six transcriptionally distinct clusters from Figure 5A, representing progenitor populations (Clusters 3 and 4), proliferating cells (Cluster 5), and effector/exhausted cells (Cluster 0). See Extended Data 5A for data from all seven clusters and for expression of additional informative genes. **(F)** Pseudobulk RNA-seq analysis of filtered (in silico) *Tcf7*⁺ TILs comparing EV and IκBδ-L OE conditions. Scatterplot of log₁₀(CPM+1) values shows genes enriched ≥2-fold in each condition. Representative (out of two independent) flow cytometric analysis of Perforin and Granzyme-B expression in OT-I TILs at day 8 following adoptive transfer of EV, IκBδ-S OE, or IκBδ-L OE cells. Histograms show TCF1 expression (*top*), and representative density plots demonstrate granzyme B⁺ perforin⁺ effector differentiation (*bottom*) within TCF1^hi^ T cells (*top*).

Pseudobulk RNA-seq analysis of *Tcf7⁺* TILs gave a quantitative measure of the increased expression of cytotoxic genes (*Gzma, Gzmb, Gzmk, and Prf1*) in this subgroup of IκBδ-L OE cells at the population level **(Figure 5F).** The *Tox*, *Nr4a1*, and *Nr4a2* mRNAs encoding exhaustion-related transcription factors were expressed at reduced levels in IκBδ-L-expressing TILs across the board, whereas the deficiency of *Nr4a3* was most evident in the Tcf7^−^ clusters (**Figure 5E-F**). At the protein level, TCF1^high^ IκBδ-L OE TILs displayed a markedly increased frequency of Granzyme B and Perforin expression compared to EV controls (**Figure 5G**), further indicating that progenitor-like cells may acquire cytotoxic function under the influence of IκBδ-L. Next, we examined the expression of specific genes in *Tcf7^+^* pretransfer cells, and in day 3 and day 8 *Tcf7^+^* TILs, and found a *Gzmb^+^Bcl6^lo^Prdm1^+^Havcr2^+^* subset that became particularly prominent in day 8 IκBδ-L expressing TILs, indicating heterogeneity within *Tcf7*^+^ TILs and accelerated differentiation into cytotoxic TILs upon IκBδ-L expression (**Extended Data 5B-G**).

### The N-terminal peptide region of IκBδ-L preferentially elicits TIL effector function

The shorter IκBδ-S isoform lacks the 143-residue N-terminal region present in IκBδ-L but retains the conserved ankyrin repeat domain (**Figure 6A**, *top*). We leveraged this naturally occurring N-terminal truncation as a functional “deletion mutant” to determine how the structure of IκBδ-L influences its proliferation and effector functions in TILs. To directly compare isoform-specific functions, we retrovirally expressed IκBδ-S or IκBδ-L in CD8⁺ T cells. Western-blotting on the day of adoptive transfer into tumor-bearing mice, four days after retroviral transduction, showed comparable expression of the two isoforms (**Figure 6A**). Bulk-RNA sequencing on the same day showed similar levels of the *Nfkbid* mRNAs (**Extended Data 6A**). The pretransfer IκBδ-L OE OT-I cells exhibited a relatively small number of differentially expressed genes compared to EV T cells (194 upregulated, 229 downregulated). Upregulated genes included activation- and effector-associated transcripts such as *Tnfrsf9* (encodes 41BB) ^39^, *Tnfrsf4* (encodes OX40) ^39^, and *Cxcl10* ^40,41^, indicating that IκBδ-L OE T cells were mildly skewed toward an effector phenotype on the day of adoptive transfer (**Extended Data 6A-C**).

**Figure 6:**
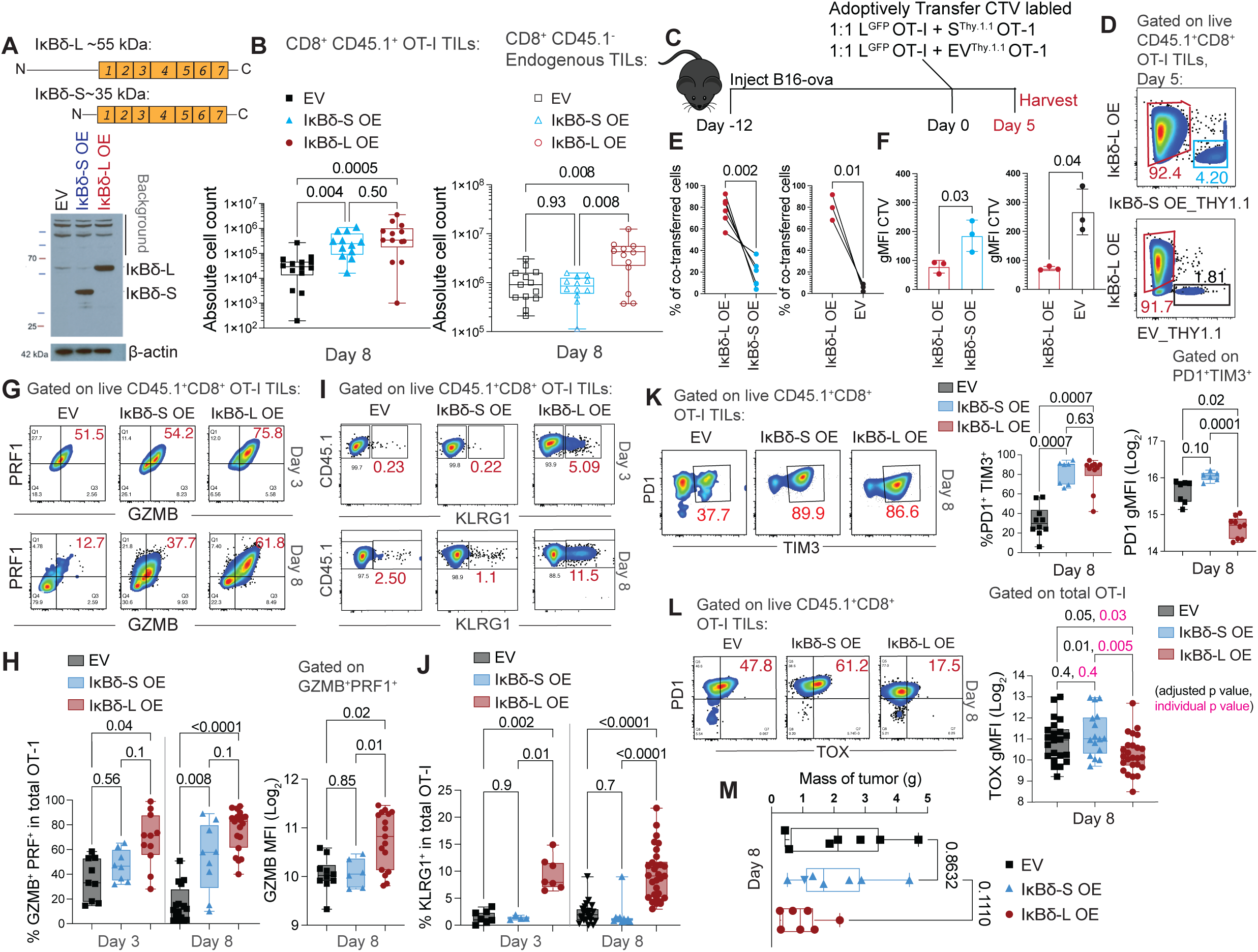
The N-terminal peptide region of IκBδ-L is necessary for acquisition of effector functions by TILs. **(A)** Schematic of IκBδ long (IκBδ-L, ∼55 kDa) and short (IκBδ-S, ∼35 kDa) isoforms illustrating the absence of a ∼150-long resisdue in the N-terminal region, and a representative western blot confirming retroviral overexpression (OE) of IκBδ-S or IκBδ-L in OT-I T cells compared with empty vector (EV); β-actin is the loading control. Data are representative of two independent experiments. **(B)** Absolute numbers of transferred CD45.1⁺ CD8⁺ OT-I tumor-infiltrating lymphocytes (TILs) (*left*) and endogenous CD45.1⁻ CD8⁺ TILs (*right*) at day 8 following B16-OVA tumor implantation, comparing EV, IκBδ-S OE, and IκBδ-L OE. Each point represents an individual mouse with n=12-14 mice per condition (OT-I: EV n=14, IκBδ-S n=12, IκBδ-L n=13; Endogenous: EV n=13, IκBδ-S n=12, IκBδ-L n=12). Statistical significance was determined using one-way ANOVA (non-parametric Kruskal-Wallis test with Benjamini and Hochberg correction for multiple comparisons for controlling the False Discovery Rate). **(C)** Experimental schematic for adoptive transfer. Mice bearing established B16-OVA tumors received a 1:1 mixture of CTV-labeled EV (THY1.1) + IκBδ-L OE (GFP), or IκBδ-S OE (THY1.1) + IκBδ-L OE (GFP) OT-I cells, followed by tumor harvest at day 5 post adoptive transfer. **(D)** Representative flow cytometry plots showing recovery of transferred OT-I populations based on GFP and Thy1.1 expression at day 5 post adoptive transfer from two independent experiments. **(E)** Percentages of co-transferred OT-I cells recovered from the same tumors, comparing IκBδ-L OE versus IκBδ-S OE (*left*) or EV (*right*) from two independent experiments. Each point represents an individual mouse with n=3-6 mice per condition. Data from two independent experiments Statistical significance was determined using paired t-test. **(F)** Geometric mean fluorescence intensity (gMFI) of CellTrace Violet (CTV) dilution in transferred OT-I cells, indicating proliferative history, comparing IκBδ-S OE and IκBδ-L OE to EV from one representative experiment with n=3 per condition. Statistical significance was determined using paired t-test. **(G)** Representative flow cytometry plots of Perforin and Granzyme-B expression in live CD45.1⁺ CD8⁺ OT-I TILs at days 3 and 8. **(H)** Quantification of the percentage of Granzyme B⁺ Perforin⁺ (GZMB⁺ PRF⁺) OT-I cells at days 3 and 8 (*left*), and granzyme B gMFI of GZMB^+^PRF1^+^ TILs at day 8 (*right*). Each point represents an individual mouse with n=9-11 for day 3, (*left*, EV n=9, IκBδ-S n=8, IκBδ-L n=11), n=9-19 for day 8 (*middle*, EV n=16, IκBδ-S n=9, IκBδ-L n=19), and n=6-17 for gMFI at day 8 (*right*, EV n=10, IκBδ-S n=6, IκBδ-L n=17). Statistical significance was determined using one-way ANOVA (non-parametric Kruskal-Wallis test with Benjamini and Hochberg correction for multiple comparisons for controlling the False Discovery Rate). **(I)** Representative flow cytometry plots showing KLRG1 expression on OT-I TILs at days 3 and 8. **(J)** Quantification of the percentage of KLRG1⁺ OT-I cells at days 3 and 8. Each point represents an individual mouse with n=4-7 for day 3 (EV n=7, IκBδ-S n=4, IκBδ-L n=7), n=10-30 for day 8 (EV n=22, IκBδ-S n=10, IκBδ-L n=30. Statistical significance was determined using one-way ANOVA (non-parametric Kruskal-Wallis test with Benjamini and Hochberg correction for multiple comparisons by controlling the False Discovery Rate). **(K)** Representative flow cytometry plots (*left*) and quantification of PD-1 and TIM-3 co-expression on OT-I TILs at day 8 (*middle*) and PD-1 gMFI in PD-1⁺TIM-3⁺ cells (*right*). Each point represents an individual mouse with n= 7-10 for frequency (EV n=10, IκBδ-S n=7, IκBδ-L n=10), and n=6-9 for gMFI (EV n=7, IκBδ-S n=6, IκBδ-L n=9). Statistical significance was determined using one-way ANOVA (non-parametric Kruskal-Wallis test with Benjamini and Hochberg correction for multiple comparisons by controlling the False Discovery Rate). **(L)** Representative flow cytometry plots for PD1 and TOX expression, and TOX gMFI in OT-I TILs at day 8. Each point represents an individual mouse with n=16-25 for Day 8 (EV n=22, IκBδ-S n=16, IκBδ-L n=25). Statistical significance was determined using one-way ANOVA (non-parametric Kruskal-Wallis test with Benjamini and Hochberg correction for multiple comparisons by controlling the False Discovery Rate). **(M)** Mass of tumors resected from mice that received either EV OT-I or IκBδ-L-overexpressing OT-I T cells. Statistical significance was determined using unpaired t-test (Day 8: EV n=7, IκBδ-S n=7, IκBδ-L n=7). All quantifications are from at least three independent experiments unless otherwise noted. All gMFI measurements were acquired using the same fluorochrome. Frequency measurements may include data from experiments employing different fluorochromes for identification of the indicated population.

We transferred 1.5 × 10⁶ IκBδ-L or IκBδ-S OE CD8⁺ T cells into C57BL/6 mice bearing established 12-day B16-OVA tumors. Tumors were harvested 8 days later to assess intra-tumoral TIL accumulation. Both IκBδ-S and IκBδ-L OE TILs accumulated to significantly higher levels than empty vector (EV) controls (**Figure 6B**, *left*). Tumors that received IκBδ-L OE T cells also had increased infiltration of endogenous host CD45.2⁺ CD8⁺ T cells (**Figure 6B**, *right*). This suggests that IκBδ-L OE TILs may actively remodel the tumor microenvironment to promote recruitment or expansion of host antitumor T cells. We performed competitive co-transfer experiments to assess the kinetics of TIL accumulation. CTV-labeled GFP⁺ IκBδ-L OE OT-I cells were mixed 1:1 with CTV-labeled Thy1.1⁺ EV or IκBδ-S OE cells and transferred into mice bearing 12-day-old tumors **(Figure 6C**). At day 5 after transfer, IκBδ-L OE TILs constituted a significantly larger fraction of recovered OT-I TILs compared to IκBδ-S or EV controls (**Figure 6D**; **Figure 6E**) and exhibited enhanced CTV dilution, indicating accelerated proliferation within the same tumor environments (**Figure 6F**).

To determine whether IκBδ-L preferentially drives effector differentiation of TILs, we adoptively transferred 1.5 × 10⁶ EV, IκBδ-S OE, or IκBδ-L OE OT-I T cells into mice bearing 12-day-old, well-established B16-OVA tumors. Tumors were harvested 3 or 8 days later, and OT-I TILs were analyzed by flow cytometry. Granzyme B and perforin are key proteins required for the cytolytic function of CD8 T cells in tumors; all TILs expressed both proteins on day 3, but IκBδ-L OE OT-I TILs contained a higher frequency of GzmB⁺Prf1⁺ TILs compared to IκBδ-S OE OT-I TILs or EV controls (**Figure 6G–H**). By day 8, IκBδ-L OE OT-I cells maintained relatively high frequencies of GzmB⁺Prf1⁺ OT-I TILs, while IκBδ-S OT-I TILs showed a modest but noticeable decrease and EV OT-I TILs showed a pronounced drop. At both early and later times in tumors, IκBδ-L OE TILs continued to display higher GZMB gMFI compared to IκBδ-S TILs (**Figure 6H**, *right*).

The observation that IκBδ-L conferred a proliferative advantage over IκBδ-S at day 5 but yielded similar total numbers of TILs at day 8 (**Figure 6B**) suggested that IκBδ-L OE TILs have a reduced capacity for long-term persistence, a phenotype characteristic of short-lived effector cells (SLECs) ^42–44^. SLECs are marked by high KLRG1 expression, an inhibitory cell surface receptor associated with terminal effector differentiation ^45^. At both days 3 and 8, IκBδ-L OE OT-I cells generated greater frequencies of KLRG1⁺ TILs than IκBδ-S OE and EV OT-I cells (**Figure 6I-J**). At day 8, both IκBδ-L and IκBδ-S OE OT-I TILs displayed elevated frequencies of PD-1⁺TIM-3⁺ cells compared to EV OT-I TILs (**Figure 6K**). However, IκBδ-L OE TILs exhibited lower levels of PD1 and TOX expression than IκBδ-S or EV TILs, consistent with an effector phenotype **(Figure 6L**). Examination of tumor sizes showed a slight, but not statistically significant, reduction in the IκBδ-L OE condition compared to the IκBδ-S OE or EV conditions (**Figure 6M**).

### IκBδ-S OE leads to the accumulation of exhausted PD1^+^TIM3^+^ TILs

To better understand the transcriptional state of IκBδ-S-overexpressing tumor-infiltrating lymphocytes, we isolated PD1⁺TIM3⁻ and PD1⁺TIM3⁺ OT-I TIL subsets from established B16-OVA tumors eight days after adoptive transfer and performed bulk RNA-sequencing (**Figure 7A**). As expected, *Nfkbid* expression was markedly enriched in IκBδ-S-overexpressing PD1⁺TIM3⁺ TILs and PD1⁺TIM3⁻ TILs (**Figure 7B**). When we compared the transcriptional profiles of IκBδ-S-overexpressing and empty-vector TILs within each subset, there were very few differentially expressed genes. Within the PD1⁺TIM3⁻ compartment, IκBδ-S OE TILs exhibited increased expression of a few genes associated with activation, proliferation, and effector differentiation, including *Batf, Irf8, Tnfsf8,* and *Cxcr6* (**Figure 7C**). Expression of genes associated with the progenitor-exhausted state was largely similar, although the level of *Sell* was slightly reduced. The expression of markers associated with proliferation ^46^, such as *Hmgb2*, *Mcm4*, *Mcm5*, *Mcm7*, *Ccna2* was also slightly reduced. The expression of exhaustion associated transcription factors, *Tox, Nr4a1, Nr4a2, and Nr4a3* were comparable. Analysis of PD1⁺TIM3⁺ TILs revealed that EV and IκBδ-S OE cells also retained comparable expression of the exhaustion-associated transcription factors *Tox, Nr4a1, Nr4a2, and Nr4a3* (**Figure 7D**). IκBδ-S-overexpressing PD1⁺TIM3⁺ TILs expressed slightly higher levels of *Entpd1* which encodes CD39, a marker of exhaustion, as well as higher levels of activation associated genes *Batf* and *Gzmk,* and lower levels of transcripts associated with killer-like cell phenotype, *Klrd1* and *Klre1*. These findings indicate that while IκBδ-S supports marked accumulation and partial activation of PD1⁺TIM3⁺ TILs, it does not repress the exhaustion-associated transcriptional network. These data further support our observation that the N-terminal region unique to IκBδ-L is required for robust cytotoxic differentiation and optimal anti-tumor effector function.

**Figure 7:**
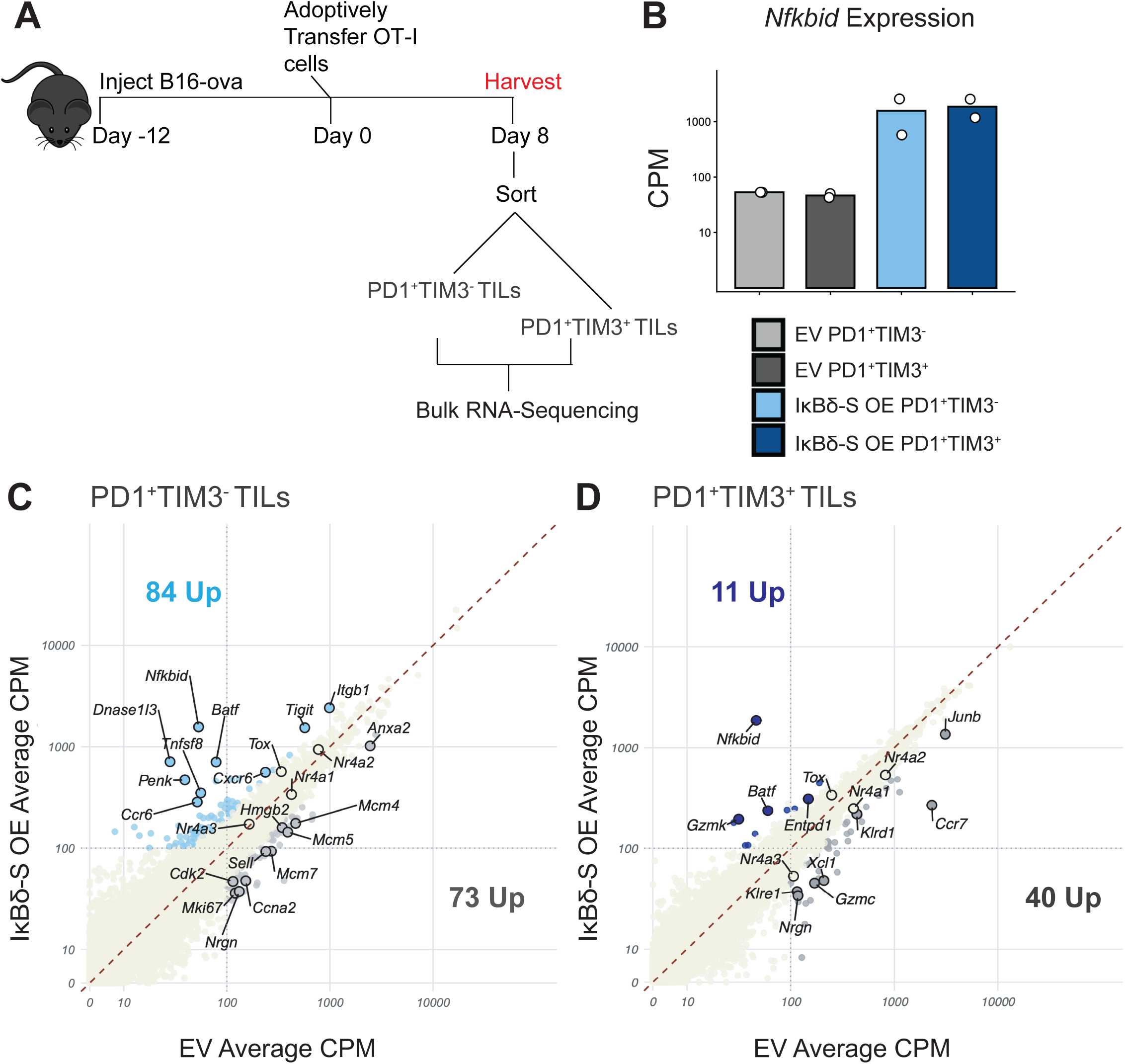
IκBδ-S OE leads to the accumulation of exhausted PD1^+^TIM3^+^ TILs without broadly reprogramming their transcriptional state. **(A)** Experimental schematic. B16-OVA tumor-bearing mice received adoptive transfer of OT-I CD8⁺ T cells transduced with empty vector (EV) or IκBδ-Short (IκBδ-S) overexpression (OE) retroviral constructs. Eight days after transfer, tumors were harvested and OT-I tumor-infiltrating lymphocytes (TILs) were FACs sorted into PD1⁺TIM3⁻ and PD1⁺TIM3⁺ populations for bulk RNA-sequencing. **(B)** Expression of *Nfkbid* in EV and IκBδ-S OE PD1⁺TIM3⁻ and PD1⁺TIM3⁺ TIL subsets. Counts per million (CPM) are shown. IκBδ-S OE TILs exhibited increased *Nfkbid* expression in both subsets, confirming transgene expression. Each symbol represents an individual biological replicate from two independent experiments. **(C)** Scatterplot comparing transcript abundance (CPM) between EV and IκBδ-S OE PD1⁺TIM3⁻ TILs averaged from individual biological replicate from two independent experiments. Genes increased in IκBδ-S OE cells are highlighted in blue and selected differentially expressed genes are annotated. Genes were highlighted based on combined thresholds of expression (≥100 CPM), statistical significance (adjusted p-value), and log2 fold-change, with selected markers annotated for visualization. The total number of genes upregulated ≥2-fold in each condition is as indicated.

### IκBδ-L OE TILs exhibit improved tumor control compared to IκBδ-S OE TILs or EV TILs

To determine whether the enhanced effector functions conferred by IκBδ-L overexpression translated to improved tumor control, we transferred 3 × 10⁶ EV, IκBδ-S OE, or IκBδ-L OE OT-I T cells into C57BL/6 mice bearing 7-day-old B16-OVA tumors (**Figure 8A**). IκBδ-L OE TILs enforced significantly better tumor control compared to both EV and IκBδ-S OE TILs during the first two weeks post-transfer (**Figure 8B**). By day 20, IκBδ-L recipients displayed significantly reduced tumor burden compared to IκBδ-S and EV controls (**Figure 8C**, *left panel*). IκBδ-S OE TILs showed a trend toward improved control relative to EV, but it did not reach statistical significance. Cumulatively, these results indicate that a large expansion of exhausted TIL numbers alone, which IκBδ-S supports, is insufficient for early tumor control. We next assessed whether improved early tumor control led to enhanced survival. Mice receiving IκBδ-L OE OT-I cells exhibited a modest extension in survival compared to EV and IκBδ-S recipient mice (**Figure 8D**). These findings are consistent with the phenotypic analyses in Figures 5 and 6 showing that IκBδ-L promotes rapid acquisition of a highly cytotoxic, KLRG1⁺ effector-like state associated with limited long-term persistence. Together, these results indicate that the combination of early expansion and effector function conferred by IκBδ-L confers improvement in early tumor control with implications for treatment of solid tumors.

**Figure 8:**
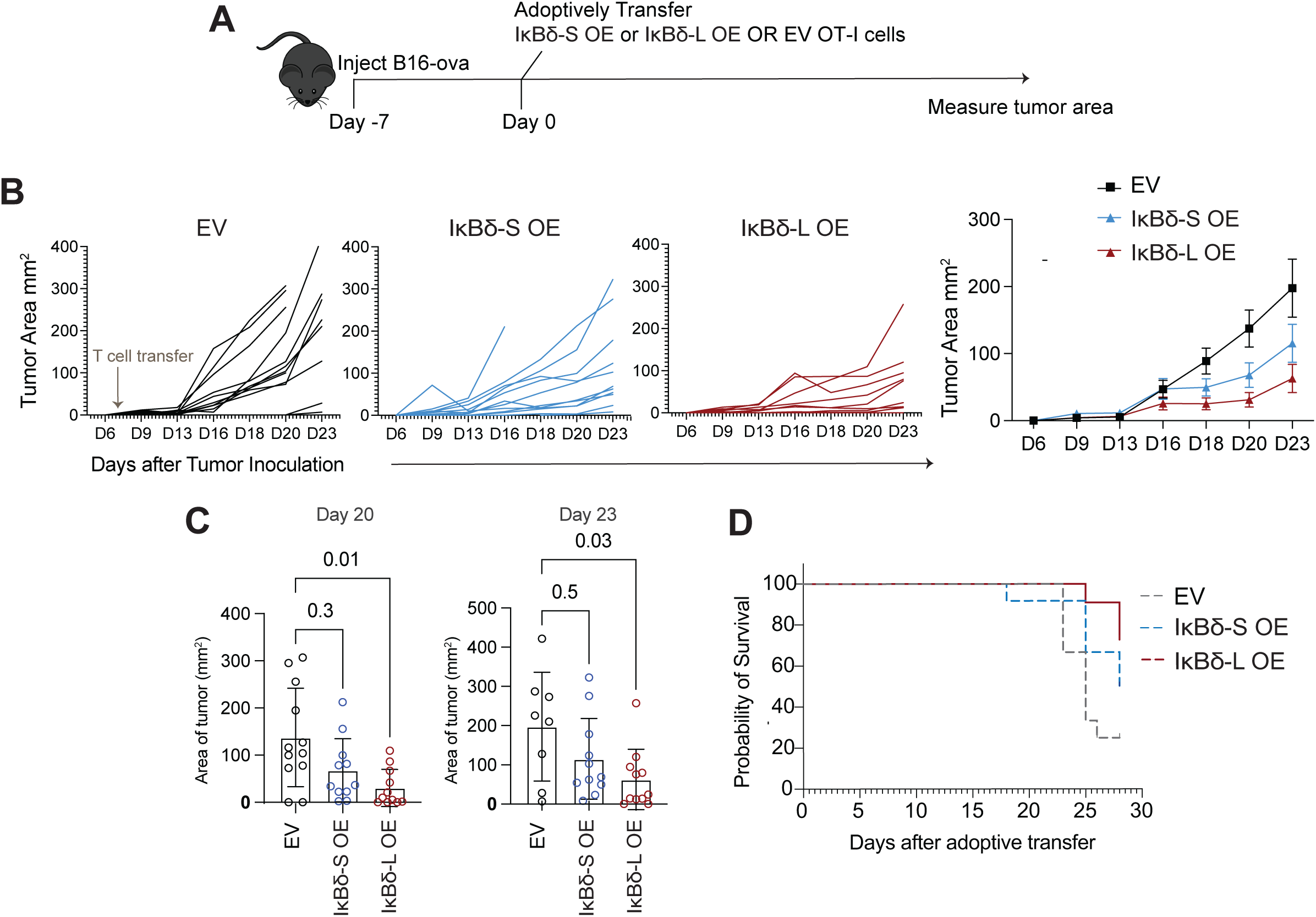
IκBδ-L OE TILs exhibit improved tumor control compared to IκBδ-S OE TILs or EV TILs. **(A)** Experimental schematic of tumor growth experiment. Mice were inoculated subcutaneously with B16 melanoma expressing ovalbumin (B16-OVA). Seven days later, mice received adoptive transfer of OT-I CD8⁺ T cells transduced with empty vector (EV), IκBδ-S or IκBδ-L overexpressing vectors. Tumor area was measured longitudinally following T cell transfer. **(B)** Individual tumor growth curves (*top*) and mean tumor area ± S.E.M. (*bottom*) over time in mice receiving EV, IκBδ-S, or IκBδ-L OT-I cells with n=11-12 (EV n=12, IκBδ-S n=12, IκBδ-L n=11). **(C)** Quantification of tumor area at day 20 with n=11-12 (*left,* EV n=12, IκBδ-S n=11, IκBδ-L n=11), and day 23 with n=8-11 (*right,* EV n=8, IκBδ-S n=11, IκBδ-L n=11) following tumor inoculation (*from* Figure 8B). Each symbol represents an individual mouse; bars indicate mean ± S.D. Statistical significance was determined using one-way ANOVA (non-parametric Kruskal-Wallis test). **(D)** Survival curves of tumor bearing mice that received adoptive transfer of OT-I CD8⁺ T cells transduced with empty vector EV, IκBδ-S or IκBδ-L expressing vectors. This is a representative experiment from 2 independent experiments.

## DISCUSSION

In this study, we have shown that the NFκB modulator IκBδ is a pivotal regulator of the anti-tumor response of CD8^+^ TILs. Overexpression of the predominant T cell isoform of IκBδ promoted proliferation and the accumulation of both PD1^+^TIM3^−^ progenitor-exhausted and PD1^+^TIM3^+^ effector-exhausted TILs, with a preferential expansion of the latter, along with sustained cytotoxic molecule expression by TILs, enhanced recruitment of endogenous CD8^+^ T cells into the tumor, and improved early tumor control. In contrast, IκBδ depletion resulted in reduced numbers of progenitor-exhausted TILs, nearly abolished the accumulation of the effector-exhausted TILs that kill tumor cells, and led to a measurable reduction in the ability of tumor antigen-specific T cells to control tumor growth. The opposite effects of IκBδ depletion and overexpression imply that IκBδ at native levels of expression in TILs contributes to CD8^+^ T cell survival/expansion and anti-tumor effector function, but that its contribution is blunted by its limited expression. IκBδ is in fact present in TILs, judging by *Nfkbid* mRNA levels, but at levels much below those of the experimentally overexpressed protein.

Our *in vitro* data cast light on the mechanisms that can be presumed to control IκBδ levels in TILs. Memory-like T cells that had been expanded in culture and rested expressed only low levels of *Nfkbid* mRNA and IκBδ-L protein at baseline, but quickly upregulated both mRNA and protein upon restimulation. Ionomycin treatment alone, mimicking the calcium branch of TCR signaling, was sufficient to evoke a modest increase in *Nfkbid* transcription, but co-stimulation with PMA was necessary for pronounced *Nfkbid* mRNA and IκBδ protein production. Thus, additional signals converge with calcium signaling for full *Nfkbid* induction. Along the same lines, direct anti-CD3 stimulation through the TCR, which elicits both calcium signaling and weak activation of MAP kinase pathways ^47^, triggered some production of IκBδ-L, which was further enhanced by activation of parallel pathways with anti-CD28. In tumors, the persistent or periodic TCR stimulation is apparently adequate to maintain some expression of IκBδ-L, but the lack of adequate co-stimulation keeps expression much below the levels that can be attained under more conducive conditions. The evidence does not yet indicate whether it is continuous expression of IκBδ or expression above a threshold level at a specific time point that directs the proliferation, survival, and— in the case of IκBδ-L— effector differentiation of TILs.

Single-cell transcriptomics demarcated clear differences between IκBδ-overexpressing TILs and controls. We found that IκBδ-L overexpression decreased the expression of exhaustion-associated transcription factors (*Tox, Nr4a1-3*) in TILs and enhanced the expression of transcripts associated with cytotoxicity (G*zma, Gzmb, Prf1, Nkg7*) as early as day 3 within the tumor. At day 8, we observed both further upregulation of transcripts that encode markers associated with short-lived effector cells (*Klrc1, Klrc2*) and a greater separation at the global transcriptional level between IκBδ-L-overexpressing TILs and empty-vector controls. Notably, we observed altered expression in IκBδ-L TILs of several genes prominent in calcium signaling— for example, *S100a6* and *S100a4* were upregulated and *Nrgn* was downregulated—raising the possibility that modulation of calcium signaling might be central to the differentiation trajectory of these cells. This is a promising area for future investigation. Closer examination of day 8 TILs documented substantial expression of *Gzmb* mRNA and GZMB protein in the progenitor-exhausted subset, which is not ordinarily characteristic of these cells. The premature expression of *Gzmb* was accompanied by reduced expression of *Myb* and *Bcl6,* and enhanced expression of *Prdm1*, indicating that IκBδ-L erodes the progenitor self-renewal program in TILs in part by accelerating acquisition of an effector program.

The functional divergence between IκBδ-L and IκBδ-S offers a point of entry for sorting out how T cells can evade the separate constraints on TIL proliferation and persistence in a tumor and on TIL effector function. IκBδ-L overexpression drove a pronounced increase in TIL numbers along with expression of the effector program. IκBδ-S overexpression led to massive accumulation of TILs without redirecting the cells along an effector pathway. These observations link the effects on TIL proliferation and survival specifically to the IκBδ ankyrin-repeat domain and its immediate flanking regions, which are shared by both IkBδ isoforms, and the effects on effector function to the N-terminal region unique to IκBδ-L. So, which protein partners interact either with both IκBδ isoforms or with IκBδ-L only? And which individual partner proteins and their corresponding downstream cellular pathways are required for TIL persistence or effector function? The answers can illuminate key choices in TIL differentiation and lead to the design of better T cells for adoptive transfer.

In summary, we report here that IκBδ-L, a newly recognized transcriptional target of NFAT, may be a useful molecular handle to steer immune responses in the tumor toward effector states in order to curb tumor growth. We had demonstrated previously that a major fraction of the chromatin regions that open in TILs in response to TCR engagement are also involved in conventional effector T cell activation, and that these regions are enriched in NFκB binding sites, thus arguably representing an underutilized “infrastructure” that could be exploited to redirect TILs toward effector states ^4^. Our new findings implement this approach in mouse tumor models by showing that overexpression of IκBδ modulates NFκB signaling and enhances early effector function in TILs. Our current study indicates that engineering CD8^+^ T cells or CAR T cells to overexpress IκBδ-L might by itself improve the response rate in human cancers where adoptive transfer therapy is already effective for some patients. An alternative strategy that is worth testing in preclinical models is combining IκBδ-L-overexpressing cells, to provide a powerful initial burst of anti-tumor function, with an additional cohort of T cells engineered to function later. Further tuning IκBδ-L expression timing or dosage, or unraveling and targeting the downstream survival and effector mechanisms controlled by IκBδ, may open new avenues to more effective and durable immune responses in the tumor.

## EXTENDED DATA

**Extended Data 1.**
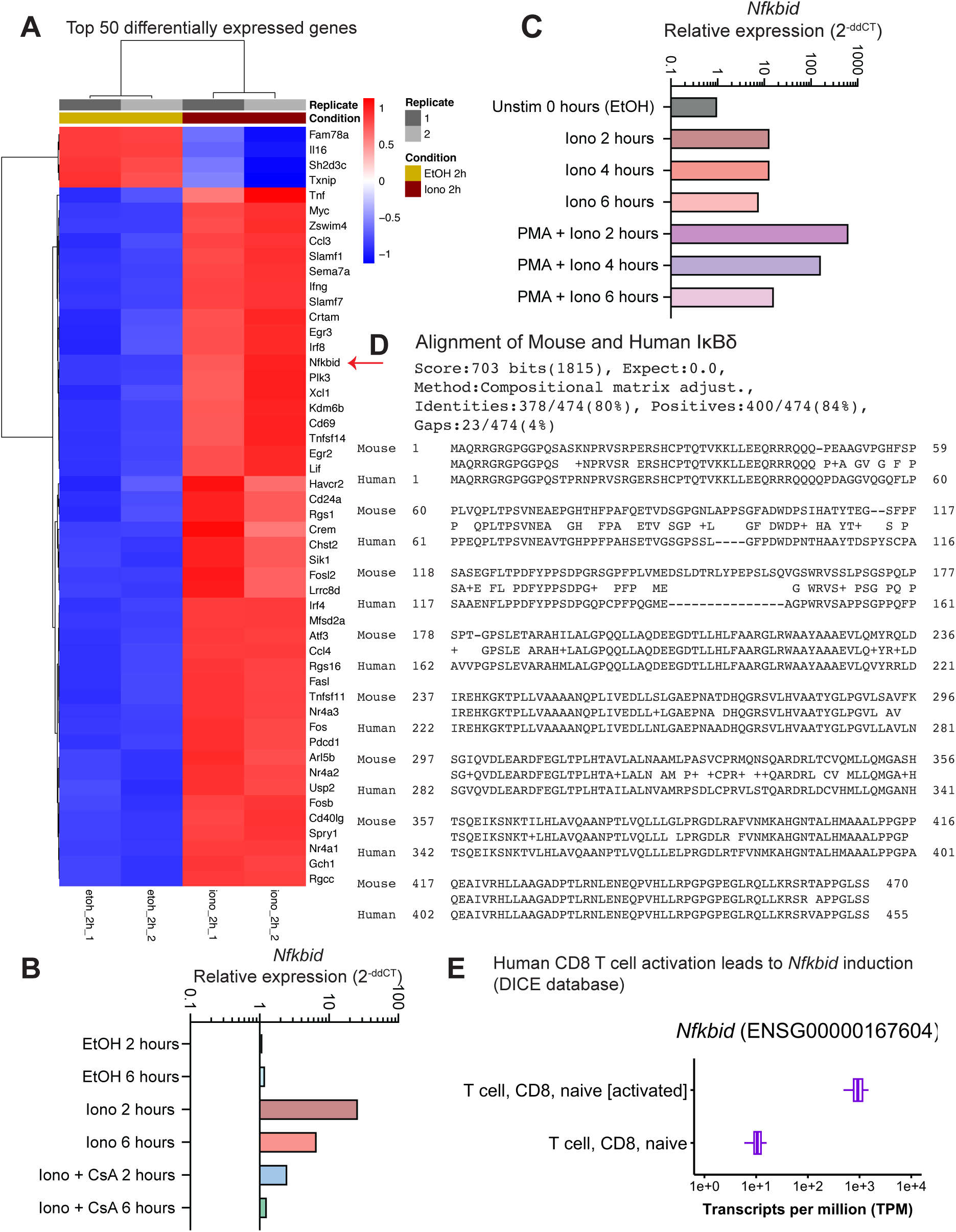
**(A)** Heatmap showing the top 50 differentially expressed genes in activated mouse CD8⁺ T cells stimulated for 2 h with ionomycin (Iono) compared with EtOH vehicle control, identified by bulk RNA-sequencing. Replicates from one representative experiment are shown. *Nfkbid* is among the significantly induced transcripts following calcium signaling and NFAT activation. **(B)** Quantitative RT-qPCR was performed to measure relative *Nfkbid* expression in unstimulated, ionomycin treated or PMA and ionomycin treated conditions at 2, 4 and 6 hours. Expression values were normalized to the housekeeping gene *Rpl13* using the ΔCq method (ΔCq = Cq*_Nfkbid_* – Cq*_Rpl13_*). Relative expression was calculated using the ΔΔCq method with 0-hour ethanol (EtOH) control sample set to 1 (2^−ΔΔCq^). Data shown are the mean of three technical replicates per condition. **(C)** Quantitative RT-qPCR was performed to measure relative *Nfkbid* expression in CD8 T cells treated with ionomycin (Iono) or EtOH ± cyclosporin A (CsA) at 2 and 6 hours to confirm bulk-RNA sequencing data shown in Figure 1C. Relative expression was calculated using the ΔΔCq method with 2-hour ethanol (EtOH) control sample set to 1 (2^−ΔΔCq^). Data shown are the mean of three technical replicates per condition. **(D)** Amino acid sequence alignment of mouse and human IκBδ proteins encoded by *Nfkbid*. Conserved residues are highlighted across the full-length proteins. Mouse and human IκBδ proteins exhibit ∼80% sequence identity and ∼84% sequence similarity. **(E)** Publicly available DICE database analysis showing induction of *Nfkbid* expression following activation of human CD8⁺ T cells, supporting conservation of activation-dependent regulation between mouse and human T cells.

**Extended Data 2.**
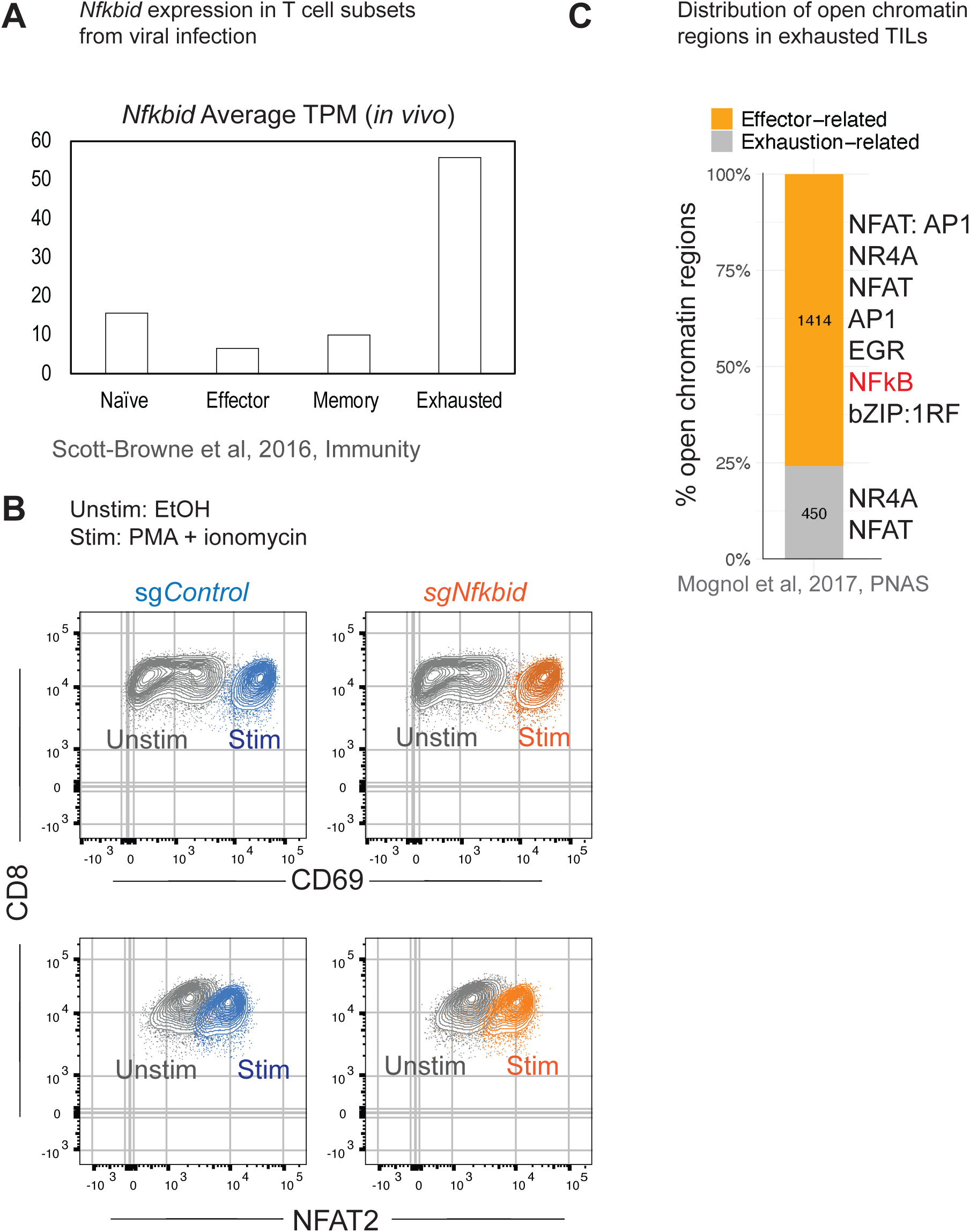
**(A)** *Nfkbid* expression (TPM) in naïve, effector, memory and exhausted T cells from acute and chronic viral infection models from Scott-Browne et al, 2016, Immunity. **(B)** Representative flow cytometry plots showing expression of CD69 and NFAT2-α in sgControl and sg*Nfkbid* OT-I CD8⁺ T cells following restimulation with PMA and ionomycin. Both sgControl and sg*Nfkbid* cells upregulated activation markers comparably, indicating comparable responsiveness despite efficient depletion of IκBδ. **(C)** Distribution of open chromatin regions previously identified in exhausted tumor-infiltrating lymphocytes (TILs), categorized as effector-related or exhaustion-related regions (adapted from Mognol et al., 2017). Effector-associated open chromatin regions are enriched for NFAT:AP-1, NFκB, AP-1, EGR, and bZIP:IRF motifs, whereas exhaustion-associated regions are enriched for NR4A and NFAT motifs.

**Extended Data 3.**
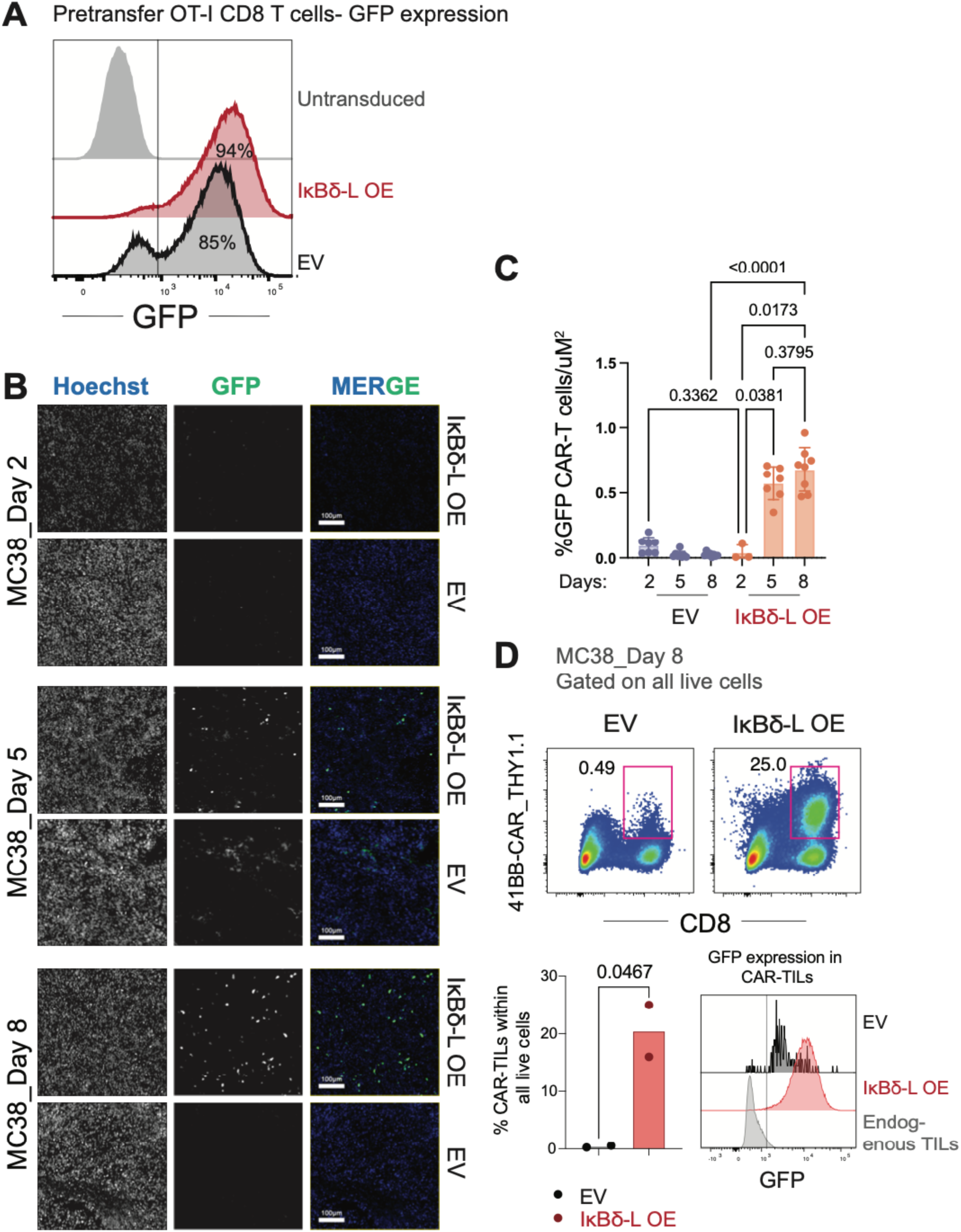
**(A)** Representative flow cytometry histograms showing GFP expression in activated CD8⁺ T cells transduced with empty vector (EV) or IκBδ-L overexpression (OE) retroviral constructs. Transduction efficiencies exceeded 85% in representative experiments. **(B)** Representative immunofluorescence images of MC38-CD19 tumors harvested at days 2, 5, and 8 following adoptive transfer of EV or IκBδ-L OE anti-CD19 CAR T cells. GFP marks transferred CAR T cells and DAPI/blue labels nuclei. IκBδ-L OE CAR T cells accumulated progressively within tumors over time, whereas EV CAR T cells remained sparse. Scale bars, 100 μm. **(C)** Quantification of GFP+ TILs/ uM^2^, each dot represents a section, representative quantification from one tumor per condition. **(D)** Confirmation of expansion of CAR-TILs in MC38-CD19 by flow cytometry in a separate experiment.

**Extended Data 4.**
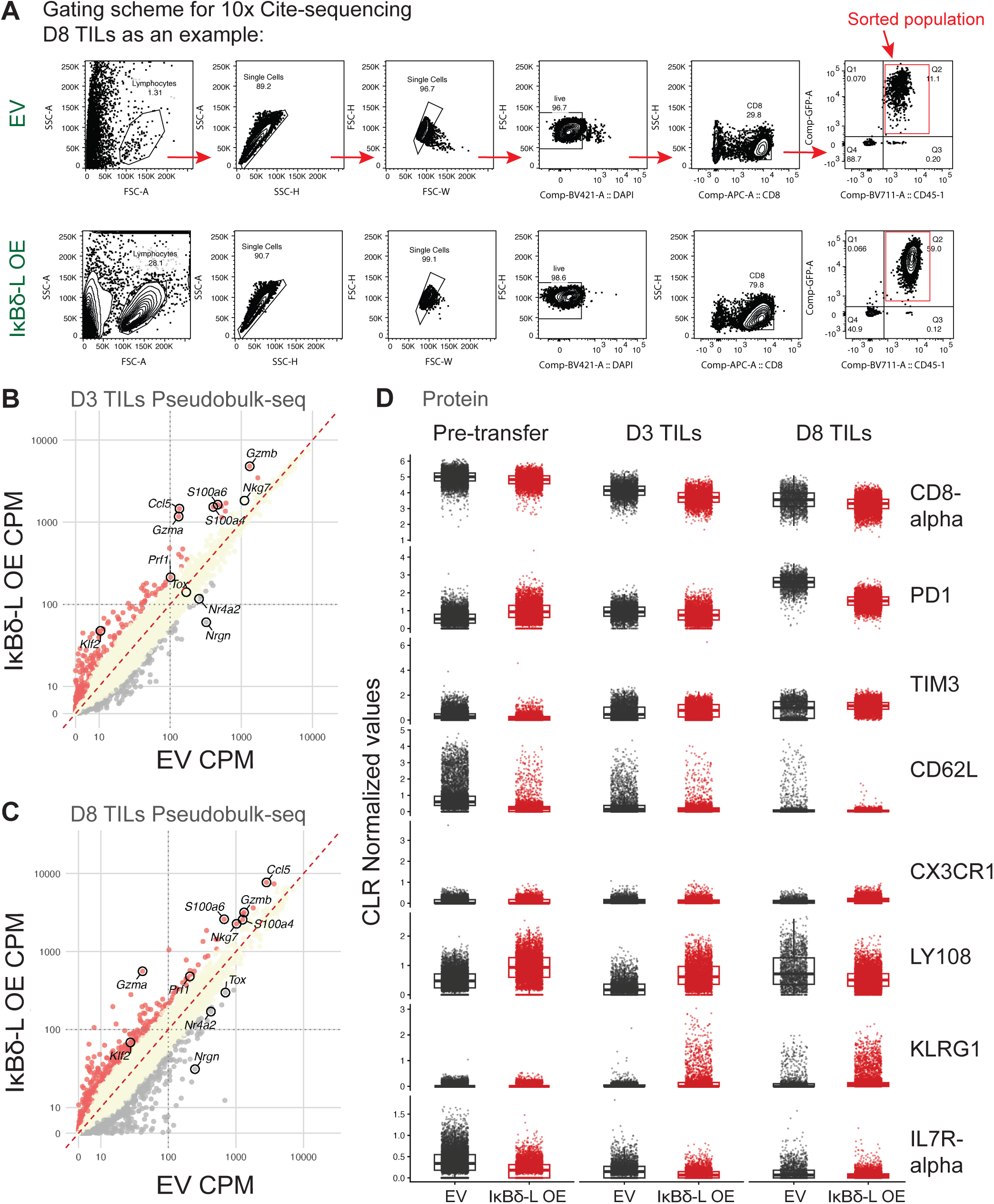
**(A)** Representative gating strategy used for sorting OT-I tumor-infiltrating lymphocytes (TILs) for 10x Genomics scRNA-seq and CITE-seq analysis. Live CD45.1⁺GFP⁺CD8⁺ OT-I cells were purified from B16-OVA tumors at day 8 after adoptive transfer of EV or IκBδ-L OE T cells. **(B)** Pseudobulk RNA-seq analysis of day 3 TILs comparing EV and IκBδ-L OE OT-I cells. Scatterplot of average CPM values highlights genes preferentially enriched in IκBδ-L OE cells, including cytotoxic molecules (*Gzmb*, *Prf1*), inflammatory mediators (*Ccl5*), and calcium-signaling regulators (*S100a4*, *S100a6*), whereas exhaustion-associated genes including *Nrgn* and *Nr4a2* were enriched in EV TILs. **(C)** Pseudobulk RNA-seq analysis of day 8 TILs comparing EV and IκBδ-L OE OT-I cells. Cytotoxic and inflammatory genes remained elevated in IκBδ-L OE TILs, whereas exhaustion-associated transcripts remained comparatively enriched in EV TILs. **(D)** CITE-seq antibody-derived tag (ADT) analysis showing CLR-normalized protein expression of CD8α, PD-1, TIM-3, CD62L, CX3CR1, LY108, KLRG1, and IL7Rα in pre-transfer OT-I cells and TILs isolated at days 3 and 8. IκBδ-L OE TILs progressively acquired a CX3CR1⁺KLRG1⁺ effector phenotype while exhibiting reduced inhibitory receptor expression relative to EV controls.

**Extended data 5.**
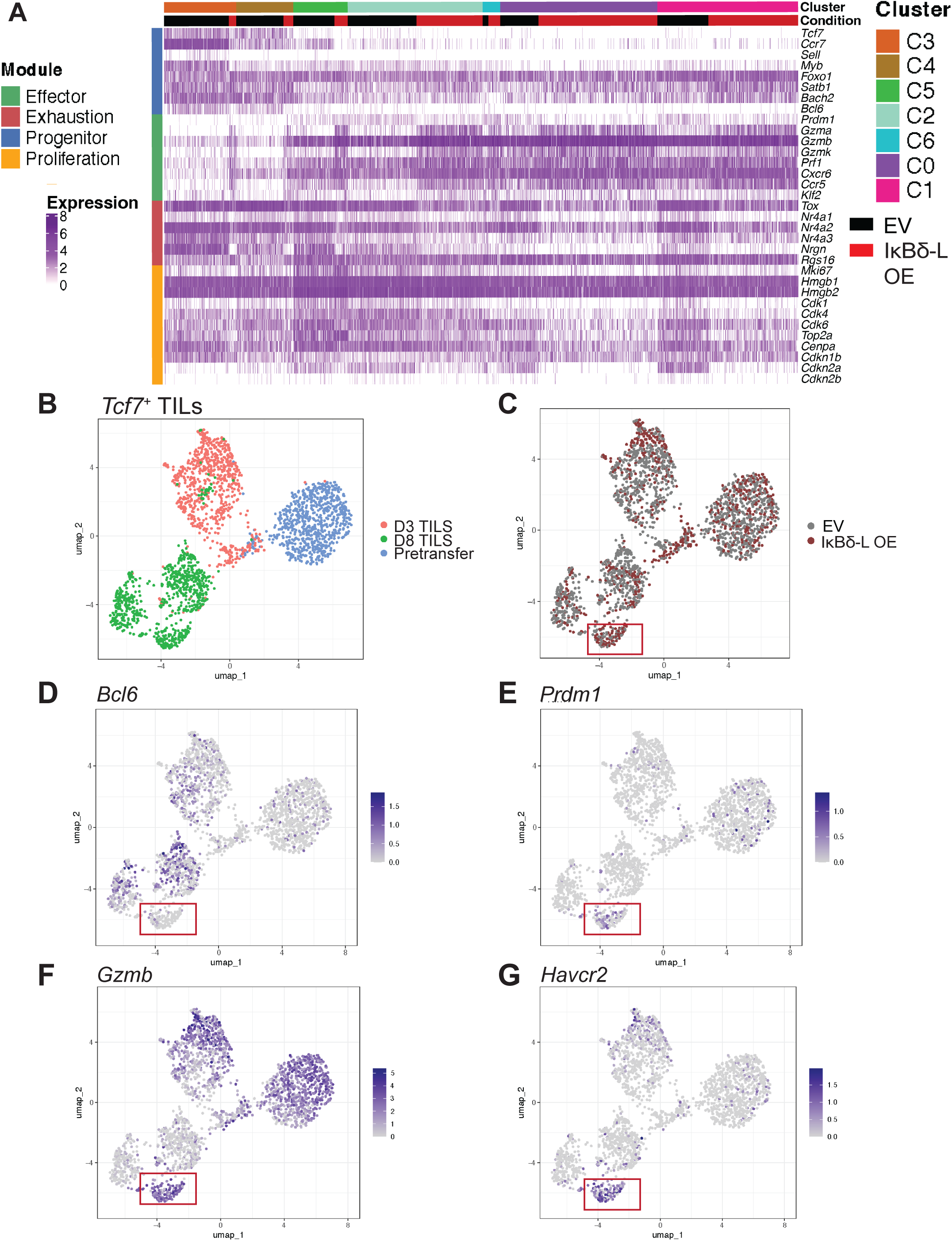
**(A)** Heatmap showing normalized expression of representative progenitor-, effector-, exhaustion-, and proliferation-associated genes across transcriptionally defined day 8 TIL clusters identified by scRNA-seq. **(B)** UMAP visualization of *Tcf7*^+^ pre-transfer OT-I cells and day 3 and day 8 TILs, illustrating progressive transcriptional divergence over time within tumors. **(C)** UMAP projection colored by EV or IκBδ-L OE condition and time. **(D–G)** Feature plots showing expression of *Bcl6*, *Prdm1*, *Gzmb*, and *Havcr2* across the UMAP space, illustrating the transition from progenitor-associated to effector-associated transcriptional states in IκBδ-L OE TILs.

**Extended Data 6.**
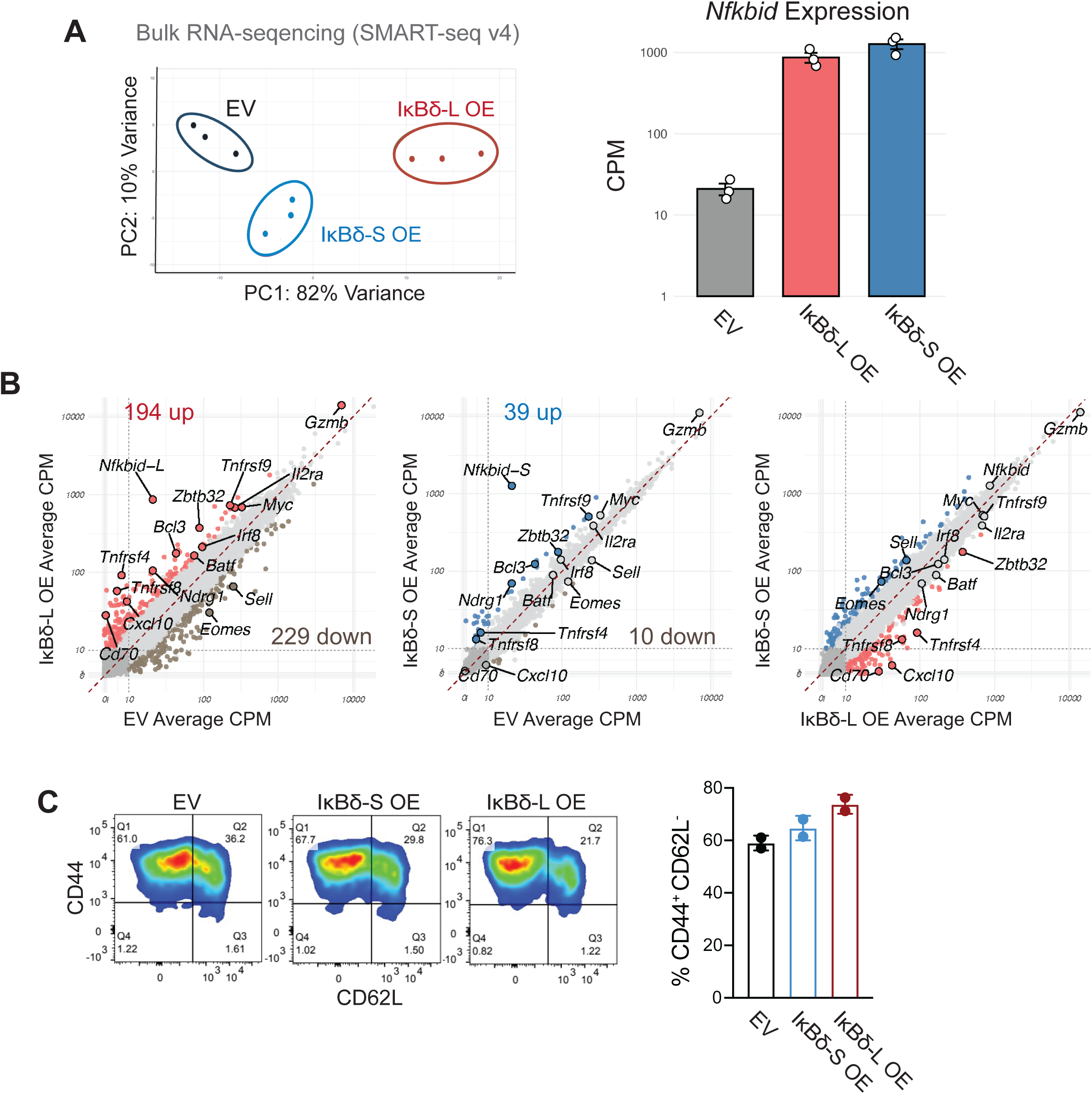
**(A)** Principal component analysis (PCA) of SMART-seq v4 bulk RNA-seq datasets from EV, IκBδ-S OE, and IκBδ-L OE OT-I CD8⁺ T cells prior to adoptive transfer showing clustering of biological replicates according to transduction condition (*left*), and *Nfkbid* expression in the three different conditions (*right*). **(B)** Scatterplots comparing pseudobulk transcript abundance between EV and IκBδ-L OE cells (left), EV and IκBδ-S OE cells (middle), and IκBδ-L OE and IκBδ-S OE cells (right). Differentially expressed genes associated with activation and effector differentiation are highlighted. **(C)** Representative flow cytometry plots showing CD44 and CD62L expression in EV, IκBδ-S OE, and IκBδ-L OE OT-I cells before transfer as well as quantifications from two independent experiments.

## MATERIALS AND METHODS

### Animals

C57BL6/J and CD45.1 (B6.SJL-PtprcaPepcb/BoyJ) mice were obtained from Jackson Laboratory. OT-I (C57BL/6-Tg (TcraTcrb)1100Mjb/J) mice with a transgenic TCR recognizing the ovalbumin (OVA257-264) peptide were a kind gift of S. Schoenberger, La Jolla Institute for Immunology, La Jolla, CA. CD45.1 OT-I mice were obtained by crossbreeding. 6-12 weeks old male and female mice were used for *in vitro* experiments. Predominantly age-matched 7-12 weeks old male mice were used as donors and recipients for *in vivo* tumor experiments. All mice were bred and maintained in the animal facility at the La Jolla Institute for Immunology under conventional specific pathogen free conditions. All experiments were performed according to protocols reviewed and approved by the La Jolla Institute Institutional Animal Care and Use Committee (IACUC) regulations.

### Cell lines and cell cultures

B16F0 mouse melanoma cell line was purchased from the American Type Culture Collection (ATCC). B16F0-humanCD19 (B16F0-hCD19) cell line was generated by transduction with amphotropic virus encoding human CD19, followed by sorting for cells expressing high levels of human CD19. The B16F10-OVA mouse melanoma cell line was kindly provided by S. Schoenberger (La Jolla Institute for Immunology). The Platinum-ERetroviral Packaging Ecotropic (PlatE) cell line was purchased from Cell Bio Labs. All tumor cell lines were tested frequently to be sure they were negative for mycoplasma contamination and were used at passage 4 after thawing from stock.

### Quantitative real-time PCR (qRT-PCR) assay

Total RNA was isolated from CD8 T cells on day 6 using Zymo Quick-RNA Miniprep Kit as per manufacturer’s protocol. qRT-PCR was performed using Power SYBR® Green RNA-to CtTM 1-step Kit (Applied biosystems) with 200ng total RNA in 20ul reaction system and analyzed on a StepOnePlus Real-Time PCR system (Applied Biosystems). The primers for *Nfkbid* (F: CCGAGGTGTTACAGATGTACCG and R:TGGTCAGTGGCGTTAGGCTCTG) and internal control *Rpl13* (F: TGAACCCAATAAAGACTGTTTGC and R: ATGTCCCCTCTACCCACAG) were ordered from IDT. The relative mRNA expression level of the *Nfkbid* was normalized to *Rpl13* expression by 2-ΔΔCT method.

### Western blotting

Cells were lysed in RIPA buffer (Thermo Fisher Scientific) supplemented with protease inhibitors (Thermo Fisher Scientific) on ice for 5-10 minutes. Lysates were clarified by centrifugation at 10,000 x g for 3 minutes at 4° C, and then treated with lambda phosphatase (New England Biolabs). For reducing conditions during electrophoresis, a solution of 1X LDS sample buffer (Thermo Fisher Scientific) supplemented with 20% ß-Mercaptoethanol was used as a loading buffer. Total protein concentration was determined using a BCA assay (Thermo Fisher Scientific). Equal amounts of protein (10-30 ug) were resolved in 4-12% NuPAGE gels and transferred to Nitrocellulose membranes (GE Healthcare Life Sciences). Membranes were blocked in 5% non-fat dry milk (Sigma) in TBS-T (Tris-HCl 15.2mM, Tris base 4.62mM and NaCl 150mM adjusted to a pH 7.5, and supplemented with 0.1% tween 20) for 1 hour at room temperature and incubated overnight at 4° C (constant shaking) with primary antibodies diluted in blocking buffer western blotting with a polyclonal Rabbit anti-IκBδ polyclonal antibody (MyBioSource). Direct-Blot™ HRP anti-β-actin Antibody (Biolegend) for loading control. After washing, membranes were incubated with HRP-conjugated secondary antibody (1:10000) for 1h at room temperature. Proteins were detected using enhanced chemiluminescence (ECL, Revvity). Membranes were exposed to CL-XPosure autoradiography film (Thermo Fisher Scientific) in a dark room and films were developed using an automatic film processor. For loading controls, membranes were washed and stripped with Restore^TM^ Western Blot Stripping buffer (Thermo Fisher Scientific, catalog # 21059) after detecting protein of interest (IκBδ isoforms) and then blotted by HRP anti-β-actin Antibody.

### Retroviral Constructs

Murine IκBδ-S/L sequences were cloned into an MSCV-based vectors that expresses GFP or Thy1.1 reporters from an IRES sequence to mark transduced cells. MSCV-based vectors encoding 41BB-CAR and Thy1.1 were also utilized. Empty vectors with appropriate reporters were utilized as negative controls.

### Transfection for retrovirus production

Transfection of plasmids was carried out with TranslT-LT1 Transfection Reagent (Mirus Bio LLC) as per manufactureŕs protocols. Briefly, 4 million Plat-E cells were seeded in T75 flasks in DMEM with 10% FBS, 1% Glutamax (Gibco), 1% penicillin/streptomycin (Gibco) 48 hours before transfection. On the day of transfection, Plat-E cells were around 80% confluent and the medium was changed once. Next, 14.25 μg retroviral plasmid and 4.75 μg pcl-Eco packaging vector plasmid were incubated with 57 μl TranslT-LT1 Transfection reagent at room temperature for 30 min in 1.9 ml Opti-MEM media (Gibco) and then carefully added to the Plat-E cells. 48 hours post transfection, virus supernatants were harvested and filtered through a 0.45 μm filter. Retro-X concentrator (Takara Bio) was used to concentrate the supernatant overnight following the manufacturer’s recommendations.

### CD8 T cell isolation and culture

Total CD8 T cells were isolated from spleens and lymph nodes of 8-12 weeks old C57BL6/J, P14 Thy1.1 or OT-I CD45.1 mice by negative selection using EasySep™ Mouse CD8^+^ T cell isolation kit (Stemcell) as per manufacturer’s protocol. In preparation for T cell activation, 6 well plates were coated with 20 μg/mL goat anti-hamster IgG crosslinking antibody (MP Biomedicals) for an hour at 37°C followed by coating with anti-CD3 (clone 2C11) plus anti-CD28 (clone 37.51) (BD Biosciences) at 1 μg/mL in home-made T cell culture medium (TCM) containing 1X MEM Essential Vitamins (Bio- Whittaker), 1X Nonessential Amino Acids (Bio-Whittaker), 1X Na Pyruvate (Bio-Whittaker), 1X Arg/Asp/Folic Acid (GIBCO, Life Technologies), 1X DME powder (GIBCO, Life Technologies) supplemented with 10% heat-inactivated fetal bovine serum, 2mM L-glutamine, 1X penicillin- streptomycin (16 units/mL Penicillin, 16 μg/mL Streptomycin), 10mM HEPES pH 7.2, and 50μM 2-mercaptoethanol. To activate CD8 T cells, a total of 3-4 million CD8 T cells were seeded at a concentration of 1 million cells/mL for 30-36 hours if cells were to undergo transduction or 48 hours for un-transduced cells. Following activation, T cells were carefully detached from the antibody-coated plate and expanded in 10 U/mL recombinant human IL-2 (PeproTech) for 4 days at 0.5 x 106 cells/mL. The concentration of T cells was adjusted to 0.5 x 106 cells/mL per day until day 6.

### Retroviral Transduction

On the day of transduction, concentrated viral supernatant was pelleted and resuspended to a concentration of 10X the original volume in TCM containing 10U/mL Rh-IL-2 and 2.6 μL/ mL Polybrene (Sigma-Aldrich) to create the transduction mix. T cells were gently removed from the activation plate at 30-36 hours post-activation and counted and replated at 4 x10^6^ per well in 6-well plates, mixed with transduction mix and centrifuged at 2000xg for 2 hours at 37°C. Cells were transduced again after 12-hour overnight rest.

### Stimulation with Ionomycin, PMA and CsA

Activated and expanded CD8 T cells were restimulated with 1μM ionomycin (Sigma), 10nM PMA (Sigma) and 1μM CsA for 2, 6 or 18 hours. All compounds were dissolved in EtOH. For unstimulated conditions, EtOH was used as vehicle only control. CsA pre-treatment was for 15 minutes prior to addition of ionomycin and/or PMA.

### Crispr/cas9 mediated knockout of *Nfkbid* in CD45.1^+^ OT-I CD8 T cells

Two crRNAs that are specific for *Nfkbid* were ordered from IDT and used separately. The DNA sequences of these two crRNAs are as follows: #1- TCGGTACATCTGTAACACCT and #2- GCTCGGGCTGTCTCCAGGGA. To prepare the crRNA-tracrRNA duplex (gRNA), 3 μL 100 μM Alt-R crRNA (IDT) and 3 μL 100 μM Alt-R tracrRNA (IDT) were mixed and annealed by heating at 95°C for 5 min, then cooled down at RT for at least 10 mins. To make the RNP mix, 4.8 μL gRNA and 2.0 uL QB3 Cas9 (80 pmol, UC Berkeley) were mixed at ∼3:1 molar ratio at RT for at least 10 mins. 1 to 2 x 106 CD8 T cells were resuspended with 25 uL electroporation buffer containing 20.4 μL P4 buffer + 4.6 μL Supplement 1 (Primary Cell 4D-Nucleofector X Kit S, Lonza). 25 uL T cells suspension and 6.8 μL RNP were mixed and incubated at RT for 2 min. The cells and RNP mix were transferred to Nucleofection cuvette strips (Primary Cell 4D-Nucleofector X Kit S, Lonza) and electroporated the cells with the 4D-Nucleofector by using the program CM137. After electroporation, the cells were cultured with TCM containing 10 U/mL human IL2.

### Alignment of human and mouse IκBδ

Protein sequences for mouse and human IκBδ were obtained from Ensembl and aligned using NCBI BLAST. Sequence similarity was assessed based on percent identity and conserved residues to evaluate cross-species conservation.

### ChIPSeq

CD8 T cells were fixed by addition of methanol-free formaldehyde (16% stock; final concentration 1%) while rotating at room temperature for 10 min. Fixation was quenched by adding 2.5 M glycine and incubating on ice for 5 min. Cells were washed three times with ice-cold PBS (2000 rpm, 5 min, 4°C) and lysed in lysis buffer supplemented with 1% Halt Protease Inhibitor Cocktail (Thermo Fisher Scientific) with constant rotation at 4°C. Lysates were centrifuged at 4,000 rpm for 5 min at 4°C, and the resulting nuclear pellet was washed sequentially with wash buffer and shearing buffer. Nuclei were resuspended in 100 μL shearing buffer, transferred to 1-mL Biorupture tube, and sonicated on a Bioruptor with 10 cycle 30S on 30S off, at 4°C, to generate chromatin fragments of the desired size(300-500bp). Debris was removed by centrifugation at 14,000 rpm for 10 min at 4°C. Chromatin concentration was quantified with the Qubit dsDNA BR Assay (Thermo Fisher Scientific). Chromatin was diluted in conversion buffer (1:9) and 5 μg of chromatin was used for each immunoprecipitation; 5% of input chromatin was reserved as an input control. Chromatin was precleared with Protein A magnetic beads (Thermo Fisher Scientific) for 1h at 4°C with rotation. Precleared material was transferred to fresh tubes and incubated overnight at 4°C with 1.5 μg anti-HA antibody (Abcam) or rabbit IgG isotype control (Abcam), together with 20 μL washed Protein A beads. Bead-bound chromatin was washed twice with RIPA buffer, once with high-salt wash buffer, once with LiCl wash buffer, and once with TE buffer, each for 10 min at 4°C with rotation. Chromatin was eluted in elution buffer for 20 min at room temperature in the presence of RNase A (0.5 mg/mL; Qiagen) with shaking at 1,000 rpm (Eppendorf Thermomixer). Protein–DNA crosslinks were reversed by incubation with Proteinase K (0.5 mg/mL final) and NaCl (200 mM final) overnight at 65°C with shaking at 1,200 rpm. DNA was purified using the Zymo ChIP DNA Clean and Concentrator Kit (Zymo Research) and quantified using the Qubit dsDNA HS Assay. Approximately 5 ng of ChIP DNA was used to generate sequencing libraries using the NEBNext Ultra II Directional RNA Library Preparation kit for Illumina (New England Biolabs), following the manufacturer’s instructions. Libraries were sequenced on an Illumina NovaSeq platform using 50 × 50 bp paired-end chemistry, targeting a depth of ∼30 million reads per sample.

### ChIPseq data processing and analysis

Paired-end ChIPseq reads were adapter- and quality-trimmed using Trim Galore (v0.6.10) and assessed with FastQC. Trimmed reads were aligned to the Mus musculus reference genome (mm10) using Bowtie2 (v2.5.3) in paired-end mode. SAM files were converted to BAM format and processed using samtools (name sorting, mate fixing, coordinate sorting, and duplicate marking). PCR duplicates were retained for peak calling with MACS2 using –keep-dup auto. Reads overlapping ENCODE mm10 blacklist regions, mitochondrial DNA, and unplaced contigs were removed using bedtools. Broad peaks were called with MACS2 (v2.2.9.1) in paired-end mode (-f BAMPE, -g mm, - -broad, --broad-cutoff 0.1, -q 0.05) using matched input controls. High-confidence peaks were defined as broadPeak entries with score >70. Peaks were annotated to genomic features using HOMER (mm10), and motif enrichment analysis was performed with findMotifsGenome.pl (default 500-bp window. Read coverage over candidate peak regions was quantified using deepTools multiBamSummary (BED-file mode) from BAM files (containing the blacklist-filtered high-confidence peaks). Genome-wide signal outside peak regions and across 10-kb bins was also quantified to assess background distribution. Raw count matrices were imported into DESeq2, and size factors were estimated based on read counts within candidate peak regions. Derived scaling coefficients were applied to generate scaled Genome-wide coverage tracks using deepTools (bamCoverage) with --normalizeUsing None and --scaleFactor.

### Tumor Experiments

Tumor cells (B16F10-OVA or B16F0-hCD19) were thawed and cultured in DMEM with 10% FBS, 1% L-glutamine, 1% penicillin/streptomycin at 37 °C in a 5% CO_2_ incubator, and were split and passaged at days 1, 3, and 5 after thawing before inoculation. On Day 0, at the time of tumor inoculation, cells were trypsinized and resuspended in Hank’s Balanced Salt Solution (HBSS, Gibco). For tumor implantation, 1.5 x10^5^ B16-OVA cells in HBSS were subcutaneously injected into the left flank of 7-12-week-old CD45.2^+^ C57BL/6J male mice. After first retroviral transduction, cells were cultured in T cell medium containing 10 U/mL recombinant human IL-2. A second transduction was performed 12 hours later using the same protocol, after which cells were cultured in T cell media containing 10 U/ml human IL-2 for 4 days. OT-I CD8 T cells were analyzed by flow cytometry to check transduction efficiency (typically greater than 90% for single retroviral transduction). On the day of adoptive transfer, mice were randomized and grouped. 1.5×10^6^ EV or IκBδ-S/L CD45.1^+^ OT-I T cells were washed and resuspended in in phosphate-buffered saline solution (PBS, Gibco) and adoptively transferred into mice bearing 12-day old tumors. Tumor-infiltrating lymphocytes (TILs) were analyzed at day 8 post adoptive transfer. To isolate the TILs, tumors were collected from the mice and placed into C tubes (Miltenyi Biotec) containing 1mg/mL collagenase D (Roche), 30 unit/mL hyaluronidase (Sigma-Aldrich) and 100 μg/mL DNase I (Sigma-Aldrich) in RPMI 1640 (Gibco) and 10% FBS. Tumors were mechanically dissociated using the gentle MACS dissociator (Milteny Biotech) followed by incubation in a shaker for 60 min at 37°C with 2000 RPM. The tumor suspension was then filtered through a 70 μM filter, and lymphocytes were separated using lymphoprep density gradient medium (STEMCELL Technologies) and stained for flow cytometry.

### Tumor growth experiments

On day 0, 7-12-week-old C57BL/6J male mice were injected subcutaneously with 2.0×10^5^ B16F10-OVA cells. When tumors were palpable, tumor measurements were recorded with a caliper 3-4 times a week and tumor size was calculated as millimeter squared (length x width). On day 7, mice were randomized and grouped and 3×10^6^ EV or IκBδ-S/L overexpressing CD45.1^+^ OT-I T cells were adoptively transferred into CD45.2^+^ C57BL/6J male tumor-bearing mice. Tumors were measured every other day for 25-30 days. For all survival experiments, tumor growth was monitored until an experimental endpoint or until IACUC-approved endpoint of a maximal tumor size measurement exceeding a diameter greater than 225 mm^2^ for more than three days without signs of regression. If mice exhibited sickness behaviors, had scars or ulcerations, adopted a hunched position, or if their body temperature was low, mice were euthanized under the guidance of the staff of the Department of Laboratory Animal Care (DLAC) at LJI. In most cases, tumor sizes were measured in a blinded manner by DLAC staff.

### Antibodies for flow cytometry

Antibodies that used for flow cytometry are as follows: anti-CD8α (clone 53-6.7, BioLegend cat# 563786, cat# 563046, cat# 100753), anti-CD45.1 (clone A20, BioLegend cat# 110739, cat# 110708), anti-PD-1 (clone RMP1-30, BioLegend cat# 109116), anti-Tim-3 (clone RMT3-23, BioLegend cat# 119723), anti-TOX (clone REA473, Miltenyi Biotec cat# 130-118-335), anti-TCF1 (clone C63D9, Cell Signaling Technology cat# 2203), anti-KLRG1 (clone 2F1, BioLegend cat# 138412), anti-CD44 (Clone IM7, BioLegend cat# 103032), anti-CD62L (clone MEL-14, BioLegend cat# 104412), anti-CD69 (clone H1.2F3, BioLegend cat# 104508), anti-NFAT2 (clone 7A6, BioLegend cat# 649605), anti-GZMB (clone GB11, BioLegend cat# 515408), anti-PRF1 (clone S16009A, BioLegend cat# 154316).

### Flow cytometry analysis

Cells were prepared in FACS buffer (PBS containing 1% FBS and 2.5 mM EDTA). Fluorochrome-conjugated antibodies used in this study were listed in table 1. All of the antibodies were used at 1:200 dilution. For surface staining, cells were resuspended in FACS buffer containing 1% mouse Fc block (Biolegend), 1:1000 diluted Fixable Viability Dye eFluor 780 (eBioscience), and stained with antibodies against surface markers for 30 min in the dark and on ice. For intracellular staining, cells were firstly stained for surface markers, briefly fixed with 4% PFA and then stained with Foxp3/transcriptional staining kit according to the manufacturer’s protocol in the dark and on ice for 30-40 minutes. Samples were measured with BD Fortessa, BD LSR III flow cytometers or Cytek Aurora, and the FACS data were analyzed by FlowJo and GraphPad Prism was used for quantitative and statistical analysis and visualization.

### The CellTrace Violet (CTV) staining

DMSO was added to a vial of CellTrace Violet (Invitrogen) staining solution to prepare a 5mM stock solution. Stock solution was diluted in PBS to generate a 5 µM working staining solution. 10 × 10⁶ in 10 mL PBS were transferred into a 50 mL conical tube, followed by centrifugation at 300 × g for 5 minutes, after which the supernatant was carefully removed. The cell pellet was then resuspended in the 5 µM CTV staining solution at the density of 10 × 10⁶ cells per mL and incubated for 20 minutes at 37 °C. Staining was quenched by the addition of 5 mL heat inactivated FBS, and the cells were incubated for an additional 10 minutes at 37 °C to allow dye efflux. Finally, the cells were centrifuged again at 300 × g for 5 minutes and washed twice with PBS and resuspended in 1.5 million cells/100 µL PBS for adoptive transfer into tumor bearing mice.

### Imaging

Immunofluorescence staining was performed on harvested mouse tumors. Tumors were isolated from mouse flanks, fixed in 4% paraformaldehyde for 48 h at 4 °C in the dark with gentle agitation. Following fixation, tissues were briefly rinsed in PBS and cryoprotected through a 5–30% sucrose gradient until fully equilibrated, indicated by tissue sinking. Samples were then embedded in Tissue-Tek® O.C.T. Compound, flash frozen on dry ice, and either stored at −80 °C or immediately sectioned. Frozen tissues were cryosectioned at 12 µm thickness, mounted onto glass slides, and stored at −80 °C until use. Prior to staining, slides were thawed at room temperature and rehydrated in PBS containing 0.1% Triton X-100 for 15 min, followed by permeabilization in PBS with 0.3% Triton X-100 for 30 min and blocking in PBS supplemented with 4% BSA for 1–2 h at room temperature. Antibodies were diluted in blocking buffer to prepare a staining master mix, which was applied to tissue sections and incubated overnight at 4 °C in a humidified chamber protected from light. After primary antibody incubation, slides were washed three times for 10 min each and subsequently incubated with secondary antibodies and Hoechst nuclear stain either overnight at 4 °C or for 2 h at room temperature. Sectioned slides were mounted using ProLong™ Diamond Antifade Mountant with #1.5 coverslips and stored in the dark at room temperature for up to three months. Sectioned slides were acquired using a slide-scanning microscope with a 20× objective. Images were analyzed in QuPath 0.6.0 for automated nuclei-based cell detection and quantification. Manual object classifiers were trained on CD45 and GFP channels to identify CD45⁺, GFP⁺, and double-positive cells within annotated tumor regions.

### Cell sorting

Cell sorting was performed by the LJI flow cytometry core, using FACS ARIA-I, FACS ARIA-II, or FACS Aria-fusion (BD Biosciences) flow cytometers using the 100uM nozzle size.

### Bulk RNA-sequencing library preparation

Cultured CD8 cells were lysed in TRIzol (Invitrogen) and then subjected to RNA isolation (ZYMO Research). To determine the gene expression changes as a result of calcium signaling with and without CsA, True-seq was utilized. For transduced T cells, The SMART-Seq v4 PLUS Kit (Takara Bio USA, Inc. Cat# R400752 & R400753) was used for library preparation as per manufacturer’s recommendations. For bulk RNA-sequencing of TILs, cells were sorted directly into lysis buffer and SMART-Seq2 protocol was used as described in PMID: 24385147. Briefly, 100 cells were collected directly into 2 μl lysis buffer (1 μl of RNase inhibitor [40U/μl, Thermo scientific] + 19 μl 0.2% (vol/vol) Triton X-100) by sorting. The SMART-Seq2 libraries were constructed following the SMART-seq2 protocol as described (literature). Libraries were prepared using the Nextera XT LibraryPrep kit (Illumina), and sequenced on an Illumina Novaseq 6000 sequencer (paired-end 50-bp reads).

### Bulk RNA-seq data processing and analysis

Paired-end Illumina RNA-seq reads were adapter-trimmed and quality-trimmed using Trim Galore (v0.6.10) with default parameters for paired-end reads, and assessed by FastQC post-trimming. Reads were aligned to the Mus musculus reference genome (mm10) using STAR (v2.7.10) in two-pass mode (allowing to map only unique reads, filtering out alignments with a ratio of mismatches/length>0.04 and specifying a minimum overhang of 8 nucleotides for novel spliced alignments to reduce false-positive splice junctions, and a minimum of 1 for pre-annotated spliced alignments to preserve sensitivity for finding known spliced junctions). Gene-level read counts were generated using HTSeq-count v2.0.4 (default settings for paired-end data) based on the UCSC RefSeq mm10 gene annotation. For purposes of visualization, coverage tracks were generated using deepTools bamCoverage (bin size = 1) and exported as BigWig files. Differential gene expression analysis was performed in R using DESeq2 (v1.42.0). Genes with < 10 counts across samples were filtered prior to modeling. Size-factor normalization was performed using the DESeq2 median-of-ratios method to account for sequencing depth and compositional differences between libraries. A generalized linear model assuming a negative binomial distribution was fit using the design formula:∼ batch + condition. This model estimated the effect of biological condition while controlling for batch as a covariate. Differential expression was assessed using Wald tests on the condition coefficient. P values were adjusted for multiple testing using the Benjamini-Hochberg procedure, and genes with a false discovery rate (FDR) < 0.05 and |FC|>2 were considered differentially expressed.

### Scatter plots for Bulk-RNA-sequencing data

RNA-seq count data (CPM) were imported and processed in R using the tidyverse framework. Gene expression scatter plots were generated using counts-per-million (CPM) normalized RNA-sequencing data. For each gene, mean CPM values were calculated across biological replicates within each experimental condition. To visualize both low- and high-abundance transcripts while preserving genes with zero expression, CPM values were transformed using a custom hybrid scale. Expression values ≤10 CPM were displayed on a linear scale, whereas values >10 CPM were displayed on a logarithmic scale. This approach maintains separation among low-expression genes while compressing highly expressed genes into a single visualization. Mean transformed CPM values were plotted for each gene, and differential expression statistics were obtained from DESeq2 analysis of raw count data. Genes were highlighted when they satisfied all of the following criteria: CPM ≥100 in at least one condition, adjusted P value (padj) <0.05, and absolute log2 fold change ≥1. Differentially expressed genes with positive log2 fold change were colored separately from genes with negative log2 fold change, while all remaining genes were displayed in gray. A diagonal reference line corresponding to equal expression between conditions and dotted lines indicating the 100-CPM threshold were included in each plot. Selected genes of interest were annotated.

### 10X Cite-seq experiment

Tumor-infiltrating lymphocytes were isolated as described above. Pre-transfer CD8⁺ T cells and processed tumor single-cell suspensions pooled from 3-4 mice per time point were incubated with fluorescently conjugated antibodies together with TotalSeq™ oligonucleotide-barcoded antibody cocktails (BioLegend) for CITE-seq profiling, including TotalSeq™-B0301 anti-mouse Hashtag 1, TotalSeq™-B0302 anti-mouse Hashtag 2, TotalSeq™-B0002 anti-mouse CD8a (Cat#100783), TotalSeq™-B0178 anti-mouse CD45.1 (Cat#110755), TotalSeq™-B0004 anti-mouse CD279 (PD-1) (Cat#109125), TotalSeq™-B0003 anti-mouse CD366 (Tim-3) (Cat#119741), TotalSeq™-B0112 anti-mouse CD62L (Cat#104465), TotalSeq™-B0563 anti-mouse CX3CR1 (Cat#149045), TotalSeq™-B0930 anti-mouse Ly108, TotalSeq™-B0250 anti-mouse/human KLRG1 (MAFA), and TotalSeq™-B0198 anti-mouse CD127 (IL-7Rα). Cells were subsequently sort-purified based on viability using Fixable Viability Dye eFluor™ 780 (ThermoFisher, 65-0865-14), CD45.1 expression (anti-mouse CD45.1 BV711, BioLegend cat# 110739), CD8α expression (anti-mouse CD8α APC, BioLegend cat# 100712), and GFP positivity. Sorting was performed using a 100-µm nozzle into collection tubes containing 100% fetal bovine serum (FBS). Collected cells were washed in PBS supplemented with 0.04% BSA and counted prior to library preparation. Single-cell RNA-seq and antibody-derived tag libraries were generated from approximately 10,000 cells per condition using the Chromium Single Cell 5′ Library & Gel Bead Kit v2 (10x Genomics) according to the manufacturer’s instructions. Libraries were pooled, quantified by the Sequencing Core at the La Jolla Institute for Immunology, and sequenced on an Illumina NovaSeq platform.

### Cite-seq filtering and analysis

Alignment, filtering, barcode counting, and UMI counting were performed using Cell Ranger v2.1.0 with default settings, and Seurat was used for further filtering. Filtering criteria included removal of cells containing excess read counts of mitochondrial genes and melanocyte genes (*Mlana*, *Dct*, *Pmel*, *Mitf*, and *Tyr*), inclusion of cells expressing *Cd8α*, and a requirement of at least 10 counts of CITE-seq CD8α antibody.

### Cite-seq data analysis and plotting

Single-cell RNA-seq data were processed using Seurat (v5.4.0). Gene expression matrices were log-normalized using the NormalizeData function, followed by identification of highly variable features with FindVariableFeatures function. Data were scaled using ScaleData, regressing out cell cycle effects based on S and G2/M scores. Principal component analysis (PCA) was performed using the top variable genes, and the first 50 principal components were used for downstream analyses. Uniform Manifold Approximation and Projection (UMAP) embeddings were generated using Harmony-corrected dimensions using the RunHarmony function. In parallel, SCTransform-based integration was performed to define cluster identities and subsequently projected onto Harmony-derived embeddings for visualization without re-clustering. This approach allowed consistent cluster annotation while visualizing condition-specific distributions in a space that preserves biological variation. Downstream analyses, including differential gene expression and visualization, were performed using Seurat and custom scripts in R using SCTransform-based clustering. Pseudobulk expression profiles were generated by aggregating raw gene-level counts across all cells within each condition (defined by vector and timepoint), thereby creating sample-level count matrices analogous to bulk RNA-seq data. For each aggregated sample, counts were normalized to counts per million (CPM) by dividing gene counts by the total library size (sum of counts across all genes) and scaling by 10⁶. Log-transformed expression values [log_10_ (CPM + 1)] were used for visualization to stabilize variance and accommodate zero values. Selected genes of interest were then extracted to compare expression patterns across experimental conditions (e.g., Pre, Day 3, Day 8) and vector groups. Violin plots and Box and Whiskers plots of protein expression were generated from CLR-normalized CITE-seq antibody-derived tag (ADT) data to visualize marker distribution across conditions. For visualization of violin plots, extreme outliers were removed by excluding values above the 99^th^ percentile within each marker, timepoint, and condition, thereby limiting the influence of rare high-expression cells on distribution shape. Gene expression between conditions was compared using scatter plots of averaged CPM values per gene, visualized using a custom hybrid transformation that applies linear scaling at low expression and log_10_ scaling at higher expression levels. Genes were highlighted exhibited equal to or greater than log2 fold-change, with selected markers annotated for visualization. Selected marker genes were visualized as heatmaps using log-normalized single-cell expression values, with cells ordered by pre-existing cluster assignments and condition. Genes were grouped into manually defined functional modules, and heatmaps were generated with cluster, condition, and module annotations. Normalized single-cell gene expression (log1p-transformed) was extracted from a Seurat object and summarized for selected genes across predefined clusters and conditions. Boxplots were generated using ggplot2 to compare expression distributions per cluster, with clusters ordered based on mean *Tcf7* expression in D8 empty vector cells.

## Acknowledgements

We would like to thank C. Kim and S. Sehic Tutusic of the Flow Cytometry Core Facility at the La Jolla Institute for immunology for cell sorting; S. Alarcon of the Sequencing Core Facility at the La Jolla Institute for immunology for next-generation sequencing; the Department of Laboratory Animal Care (DLAC) and members of the animal facility for expert animal support, especially Fernando Vazquez; Roberta Nowak, project manager at the Division of Signaling and Gene Expression, La Jolla Institute for immunology, for her excellent support; and all members of the Rao, Hogan and Myers labs for insightful comments and discussion.

## Author contributions

BD, XH, and PGH designed the study. BD, XH, AL-C, YZhao, HD, OI, EJ, YZhu, CZ, and SB performed experiments. BD, AL-C, LJA-V, and EG-A carried out bioinformatic analyses. BD, AR, and PGH analyzed data with input from the other authors. BD and PGH wrote the manuscript. BD, AR and PGH edited the manuscript.

## Data and materials availability

RNA sequencing data is available in the BioProject database under the accession number PRJNA1478066. Additional information and material related to this manuscript will be available upon request.

